# Controls on the isotopic composition of microbial methane

**DOI:** 10.1101/2021.09.14.460204

**Authors:** Jonathan Gropp, Qusheng Jin, Itay Halevy

**Affiliations:** Department of Earth and Planetary Sciences, Weizmann Institute of Science, Rehovot, Israel; Department of Earth Sciences, University of Oregon, Eugene, Oregon, USA

## Abstract

Microbial methane production (methanogenesis) is responsible for more than half of the annual emission of this major greenhouse gas to the atmosphere. Though the stable isotopic composition of methane is often used to characterize its sources and sinks, strictly empirical descriptions of the isotopic signature of methanogenesis currently limit such attempts. We developed a biochemical-isotopic model of methanogenesis by CO_2_ reduction, which predicts carbon and hydrogen isotopic fractionations, and clumped isotopologue distributions, as functions of the cell’s environment. We mechanistically explain multiple-isotopic patterns in laboratory and natural settings and show that such patterns constrain the in-situ energetics of methanogenesis. Combining our model with environmental data, we infer that in almost all marine environments and gas deposits, energy-limited methanogenesis operates close to chemical and isotopic equilibrium.

## Introduction

Methane (CH_4_) is a major greenhouse gas, with both natural and anthropogenic sources (1). The primary natural source of biogenic methane emissions is archaeal methanogenesis in anoxic environments (2), about a third of which is hydrogenotrophic (reduction of CO_2_ with dihydrogen, H_2_; Ref. 3). Strong isotopic discrimination during biological and abiotic methane formation has motivated the use of methane hydrogen and carbon isotopes to trace its production and consumption processes, construct global methane budgets and evaluate its climatic impacts (1, 4, 5). Current organism-level models that rely on isotopic mass balance can explain part of the observed range of microbial isotopic discrimination (6–9), but to date, such models have prescribed rather than resolved the microbial biochemistry. It has been difficult, therefore, to distinguish between different methane sources, different modes and extents of environmental methane cycling, and different environmental controls on the microbial isotope discrimination as drivers of observed variations in the isotopic composition of methane. To constrain the microbial component of such variations, we developed and analyzed a full metabolic-isotopic model of hydrogenotrophic methanogenesis with the net reaction:

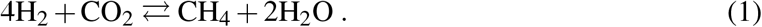

The model predicts the isotopic discrimination and its relation to the thermodynamic drive of this pathway (the Gibbs free energy of the net reaction, ΔG_net_) and to cell-specific methanogenesis rates in laboratory cultures. Extending our analysis to energy-limited conditions, which are prevalent in natural environments, our model reveals the environmental and metabolic controls on the isotopic composition of methane.

## Results and Discussion

### A metabolic model of hydrogenotrophic methanogenesis

Accounting for the kinetics and thermodynamics of enzymatically-catalyzed reactions in hydrogenotrophic methanogenesis (details in the Methods and Supplementary Materials, SM), we constructed mass balance equations for the intracellular metabolites in the pathway (Fig. 1A). Given extracellular concentrations of CO_2_, H_2_ and CH_4_, and pH, which define ΔG_net_, these equations are solved for the steady-state concentrations of the intracellular metabolites (fig. S1). At this steady state, the model links metabolite concentrations to the net rate of methanogenesis, and to the gross forward and reverse rates of the individual reactions. We calibrated our metabolic model to available measurements of specific methanogenesis rates and their relation to H_2_ concentrations in the *µ*M to mM range (fig. S2; SM). The metabolic model accounts for electron transfer from H_2_ to the intermediate metabolites through three electron carriers, cofactor F_420_, coenzyme B (HS-CoB), and ferredoxin (Fd), thereby providing insight into the oxidation state of the cell. We validated the model’s predictions for the oxidation state of these electron carriers against available measurements. Fmd catalyzes the first step in the pathway, CO_2_ fixation to organic carbon with two electrons from an iron-sulfur-containing Fd (10). We found that to drive this reaction in the direction of net methanogenesis, Fd needs to be *>*90% reduced, and that with decreasing CO_2_ concentrations, Fd must approach 100% reduced to compensate for the decreasing thermodynamic drive (fig. S1J). This is in line with previous estimates of the oxidation state of Fd for this reaction (11). F_420_ is 0.1% reduced at low H_2_ concentrations, and up to 100% reduced at high H_2_ concentrations (fig. S1K), in agreement with observations from lab cultures (12). Coenzyme B (HS-CoB) is 10% reduced at low H_2_ concentrations, and up to 100% reduced at high H_2_ concentrations (fig. S1L). The lower range is in agreement with observations of the oxidation state of HS-CoB at *µ*M to 1 mM H_2_ concentrations (13). Our observation-validated model predictions suggest that the intracellular oxidation state of electron carriers generally does not reflect electrochemical equilibrium, and is instead maintained dynamically, by oxidation and reduction fluxes that depend on the cell’s metabolic activity. This dynamic control means that measurements of the oxidation state of specific reduction-oxidation couples does not necessarily bear on the oxidation state of other reduction-oxidation couples (both inorganic and organic) and on the overall oxidation state of the cell.

**Figure 1:**
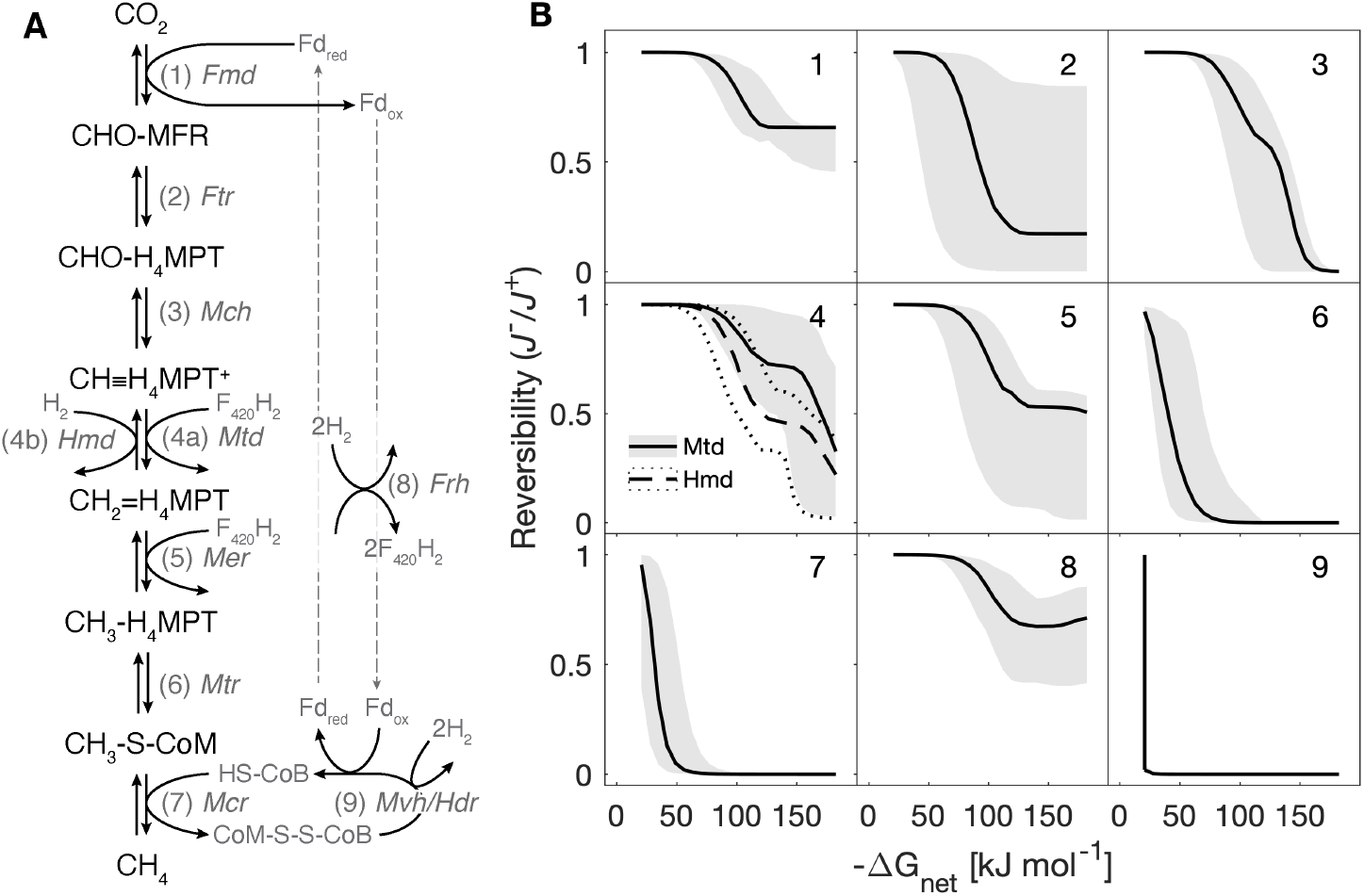
A metabolic model of hydrogenotrophic methanogenesis. (**A**) Schematic pathway. Abbreviated enzyme names are in italics (table S1). (**B**) Individual reaction reversibility 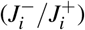 against ΔG_net_. The subplots in panel B are numbered in accordance with panel A and table S1. The black lines are the median of 10^3^ model simulations, and envelopes represent 95% of model results. The uncertainty originates from the enzyme kinetic parameters. Simulations were carried out at [H_2_] of 1 nM to 10 mM, and typical experimental [CO_2_] and [CH_4_] of 10 mM and 10 *µ*M, respectively.

The calibrated and validated biochemical model forms the basis for the metabolic-isotopic coupling. With increasing H_2_ concentrations, ΔG_net_ becomes increasingly negative, and the individual reactions in the pathway depart from equilibrium to various extents (Fig. 1B). We quantify the departure from equilibrium by the individual reaction reversibility, defined as the ratio of the reverse to forward gross rates of that reaction and related to its actual transformed Gibbs free energy 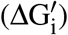 (14–16):

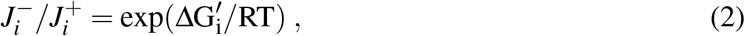

where R is the gas constant (J mol^−1^ K^−1^), T is the temperature (K) and 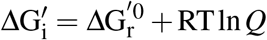 (J mol^−1^, where *Q* is the reaction quotient). For net forward methanogenesis rates, the reversibility of individual reactions varies between zero (a near-unidirectional, kinetically controlled reaction) and unity (a near-equilibrium reaction). Some reactions do not fully depart from reversibility in the explored ΔG_net_ space (e.g., the Frh-catalyzed reduction of F_420_), whereas others depart from reversibility at ΔG_net_ as modestly negative as –15 kJmol^−1^ (e.g., the Mvh/Hdr, Mcr and Mtr-catalyzed reactions). This differential response is a combined function of the reaction thermodynamics, expressed as the standard-state transformed Gibbs free energy 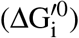, and the enzyme kinetics, specifically the metabolic rate capacity (*V* ^+^) and Michaelis constants (*K*_*M*_). Neither 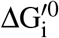 nor *V* ^+^ and *K*_*M*_ in isolation predict the pattern of differential departure from equilibrium—coupled thermodynamics and kinetics must be considered.

### An isotopic model calibrated to laboratory cultures

The isotopic discrimination between substrate (*s*) and product (*p*) is described by the isotopic fractionation factor ^*r*^*α*_*s/p*_ = ^*r*^R_*s*_*/*^*r*^R_*p*_, where ^13^R = ^13^C/^12^C, ^2^R = D/H (with D ≡ ^2^H and H ≡ ^1^H), and “*r*” denotes the rare isotope. The net isotopic fractionation expressed in an individual (bio)chemical reaction may vary between thermodynamic equilibrium and kinetic end-members, associated respectively with a reversible reaction and unidirectional forward reaction to form the reaction product. With the reversibilities calculated in the metabolic model and with values assigned to the equilibrium and kinetic isotopic fractionation factors (EFFs and KFFs, respectively) of the individual reactions, the net isotopic fractionation between the pathway substrates and products may be calculated (17, 18). We calibrated the relations between ΔG_net_ and the resulting net carbon and hydrogen isotopic fractionations against experimental data, then used the calibrated model to determine the relation between ΔG_net_ and the abundance of doubly substituted (“clumped”) methane isotopologues, which has yet to be systematically explored in experiments.

### Bulk carbon and hydrogen isotopic fractionation

In laboratory cultures, the CO_2_–CH_4_ carbon isotope fractionation (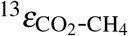, where *ε* = 1 − *α* [‰]) is inversely related to ΔG_net_ (8, 19–23, fig. S3A). At near-zero ΔG_net_, the individual reactions in the pathway operate close to equilibrium 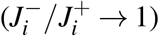, and our model predicts 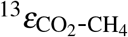 close to the temperature-dependent isotopic equilibrium fractionation 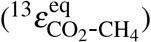. When ΔG_net_ becomes slightly negative, 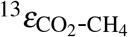 peaks to larger-than-equilibrium values of 80-100‰ at ≈–45 kJmol^−1^, followed by a gradual decline to ≈30‰ reached at ΔG_net_ of ≈–120 kJmol^−1^ (Fig. 2A). Such larger-than-equilibrium 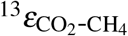 values have been observed in several experimental and environmental datasets and have yet to be explained mechanistically. Our model reveals that this 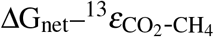 relation is controlled by the landscape of departure from equilibrium of the individual reactions in the pathway. The carbon reaction network in methanogenesis is linear. In such networks, near-unidirectionality of an individual reaction 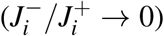 leads to expression of that reaction’s KFF and suppresses isotopic fractionation associated with downstream reactions (18). The Mcr-catalyzed reaction’s KFF (≈40‰, Ref. 24) is larger than its EFF (≈1‰, Ref. 25). As this reaction departs from equilibrium, the peak in 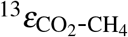 reflects a sum of its KFF and the EFFs of upstream reactions, with the exception of the Mtr-catalyzed reaction, which also partially departs from equilibrium (Fig. 1B). The result is a larger-than-equilibrium 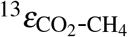 at modestly negative ΔG_net_ values. The 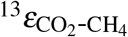 floor at ΔG_net_ more negative than ≈–120 kJmol^−1^ (Fig. 2A) is defined by partial expression of the KFFs of Ftr and Fmd, and suppression of the isotopic fractionations associated with downstream reactions.

**Figure 2:**
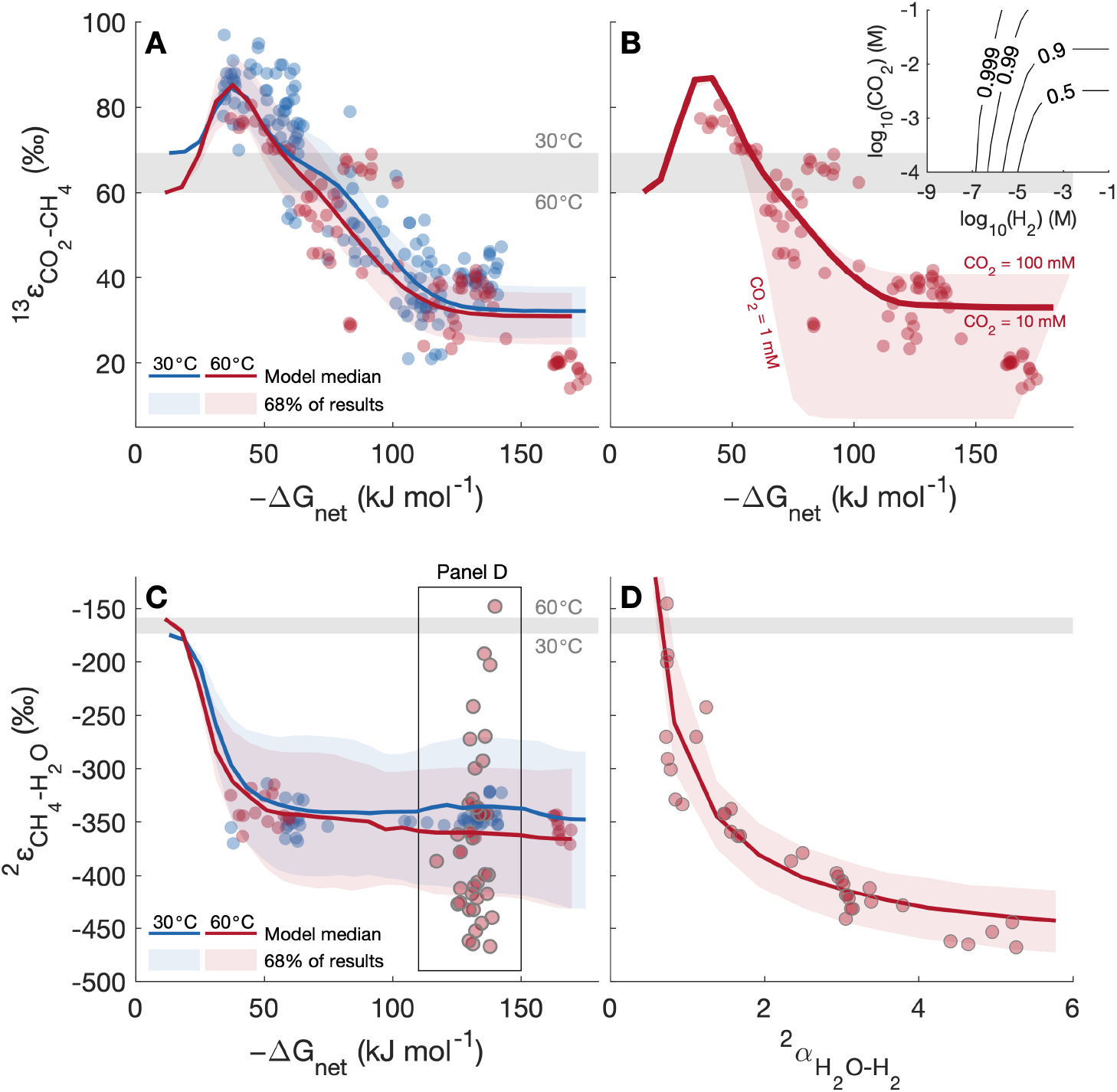
Model-laboratory culture comparison of bulk carbon and hydrogen isotopic fractionation. Mesophilic (30–40°C, blue circles) and thermophilic (≥55°C, red circles) experimental data and model results (lines and envelopes) at 30°C (blue) and 60°C (red), and the equilibrium isotopic fractionation (gray envelopes). (**A, C**) 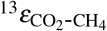 and 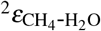 against ΔG_net_. (**B**) 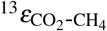 against extracellular CO_2_ concentrations ([CO_2(out)_]). Contours in the inset show the reversibility of cross-membrane CO_2_ diffusion. (**D**) Mixing effects on 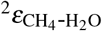 as a function of the H_2_O–H_2_ isotopic fractionation 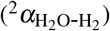, compared to laboratory culture data (8).

At large-negative ΔG_net_ values 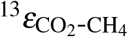 is sensitive also to the extracellular partial pressure of CO_2_ (*p*CO_2_). At a steady state, intracellular CO_2_ utilization is exactly matched by net CO_2_ diffusion across the membrane. This net diffusive flux is the difference between large gross fluxes (into and out of the cell) when *p*CO_2_ is high, and smaller gross fluxes at lower *p*CO_2_. Thus, the reversibility of net diffusion is low at low *p*CO_2_, and suppression of downstream net carbon isotopic fractionation results in small 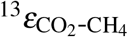 (also referred to as a “reservoir effect”). The dependence of 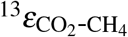 on *p*CO_2_ explains the smallest net fractionations observed in laboratory cultures (lower bound of red envelope in Fig. 2B, Ref. 21), as well as the dependence of 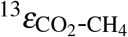 on pH in hyperalkaline settings (26, 27).

The CH_4_–H_2_O hydrogen isotopic fractionation 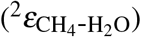 in laboratory cultures is ≈200‰ more negative than the temperature-dependent isotopic equilibrium fractionation 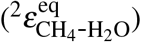, and existing observations suggest that it does not display a clear dependence on ΔG_net_ (8, 19, 22, 28, 29, fig. S3B). Unlike the linear carbon reaction network, the hydrogen reaction network has four branches, each of which has the potential for hydrogen atom exchange between pathway intermediates and H_2_O. Therefore, departure from equilibrium of one of the hydrogen atom exchange reactions does not preclude CH_4_-H_2_O hydrogen isotopic equilibrium. Specifically, the Mvh/Hdr-catalyzed reaction is near-irreversible at ΔG_net_ values as high as –25 kJmol^−1^ (Fig. 1B), yet at this ΔG_net_ value 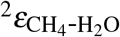 is approximately equal to 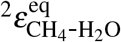 (Fig. 2C), and this arises from the high reversibility of the other hydrogen atom exchange reactions in the pathway. Only when the Mcr- and Mtr-catalyzed reactions sufficiently depart from reversibility (at ΔG_net_ ≤ –30 kJmol^−1^), cutting off CH_4_ and HS-CoB from exchange with upstream intermediates that are close to hydrogen isotope equilibrium with H_2_O, does 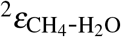 depart from 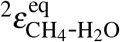 (Fig. 2C). At ΔG_net_ more neg ative than −40 kJmol^−1^ the KFFs of Mcr, Mtr and Mvh/Hdr control 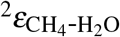 (fig. S4). Overall, our model reveals a clear 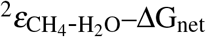 relation, which has not been accessed by the range of ΔG_net_ explored in laboratory cultures to date. The apparent invariance of 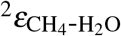 at ΔG_net_ more negative than –40 kJmol^−1^ represents a complex, ΔG_net_-dependent combination of isotope effects associated with the enzymes in the pathway.

The hydrogen atom added to CH-H_4_MPT may come from H_2_O in the Mtd-catalyzed reaction, or from H_2_ in the Hmd-catalyzed reaction (Fig. 1A; 30). Thus, up to one quarter of the hydrogen atoms in methane may come from H_2_, depending on the relative activity of Mtd and Hmd. High H_2_ concentrations favor high methanogenesis rates and Hmd activity. Under these conditions, and especially in the case of H_2_-H_2_O isotopic disequilibrium, 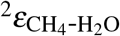 may vary in response to variations in the hydrogen isotopic composition of H_2_. Our model captures this behavior, as observed in culture experiments with 1 mM [H_2_] and at 60°C (8, 29), displaying 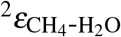 values between –145‰ and –480‰, which inversely covary with the H_2_O-H_2_ isotopic fractionation (Fig. 2D).

Our predicted trajectories for departure from equilibrium 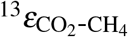 and 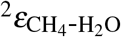 values may explain observations of near-equilibrium 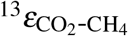 values concurrent with clearly disequilibrium 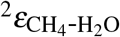 values, which have been previously explained by a decoupling of the carbon and hydrogen isotopic systems in methanogenesis (8, 31). We suggest instead, that the measured 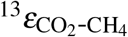 values did not reflect isotopic equilibrium, but the descending branch from the 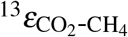 maximum (which occurs at ΔG_net_ ≈–40 kJmol^−1^) with increasingly negative ΔG_net_. In other words, we suggest that apparent equilibrium 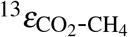 values may emerge by a fortuitous combination of EFFs and KFFs, and not due to actual isotopic equilibrium between CO_2_ and CH_4_.

### Clumped isotopologue distributions

The abundances of the doubly substituted isotopologues of CH_4_ (^13^CH_3_D and ^12^CH_2_D_2_) are expressed as a deviation from their concentrations at a stochastic distribution of the rare isotopes, with Δ^13^CH_3_D = ^13,2^*R*_sample_*/*^13,2^*R*_stochastic_ −1 [‰] and Δ^12^CH_2_D_2_ = ^2,2^*R*_sample_*/*^2,2^*R*_stochastic_ − 1 [‰]. Both Δ^13^CH_3_D and Δ^12^CH_2_D_2_ values depend on the methane formation temperature (6), yet applications of methane clumped isotopes to constrain its formation temperature and mechanism are complicated by source mixing and disequilibrium effects (7, 32). The dependence of Δ^13^CH_3_D and Δ^12^CH_2_D_2_ on ΔG_net_ has not been determined experimentally, and similar to bulk carbon isotopes, our model predicts non-monotonic departure from clumped isotopic equilibrium (fig. S5), unlike previous estimates of these relations (7, 33). As ΔG_net_ becomes negative, both Δ^13^CH_3_D and Δ^12^CH_2_D_2_ values decrease from the expected equilibrium compositions, and Δ^12^CH_2_D_2_ becomes anti-clumped (i.e., *<* 0‰) due to expression of the KFFs of the Mcr- and Mtr-catalyzed reactions (fig. S5). After the initial decrease in Δ^13^CH_3_D and Δ^12^CH_2_D_2_, both increase with increasingly negative ΔG_net_, and Δ^13^CH_3_D increases to values almost as high as the equilibrium values. This behavior has two implications. First, there is a range of ΔG_net_ (≈–75 to –55 kJmol^−1^) over which Δ^13^CH_3_D values may give the false appearance of proximity to isotopic equilibrium. Second, there is a range of ΔG_net_ (≈–100 to –20 kJmol^−1^) over which Δ^13^CH_3_D and Δ^12^CH_2_D_2_ cannot uniquely constrain the energetic state of the cell (e.g., Δ^13^CH_3_D is ≈4 ‰ at both ΔG_net_ of ≈–20 and ≈–70 kJmol^−1^). However, in combination with 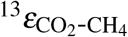 and 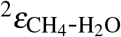 data, the position in the ΔG_net_ landscape and the degree of departure from equilibrium may be uniquely constrained (Fig. 3). The multiple-isotope composition of methane (i.e., bulk carbon and hydrogen, and clumped isotopes) may thus be a useful proxy for ΔG_net_ in natural environments where measurements of H_2_, CO_2_ and CH_4_ concentrations are not easily obtainable. In addition to departure from equilibrium of the Mcr- and Mtr-catalyzed reactions, which may cause anti-clumped Δ^12^CH_2_D_2_ compositions, a testable prediction of our model is that the Hmd-catalyzed reaction may also cause anti-clumping. This arises from combinatorial effects (34–36) when Hmd activity is high, and especially during H_2_-H_2_O hydrogen isotopic disequilibrium (fig. S6).

**Figure 3:**
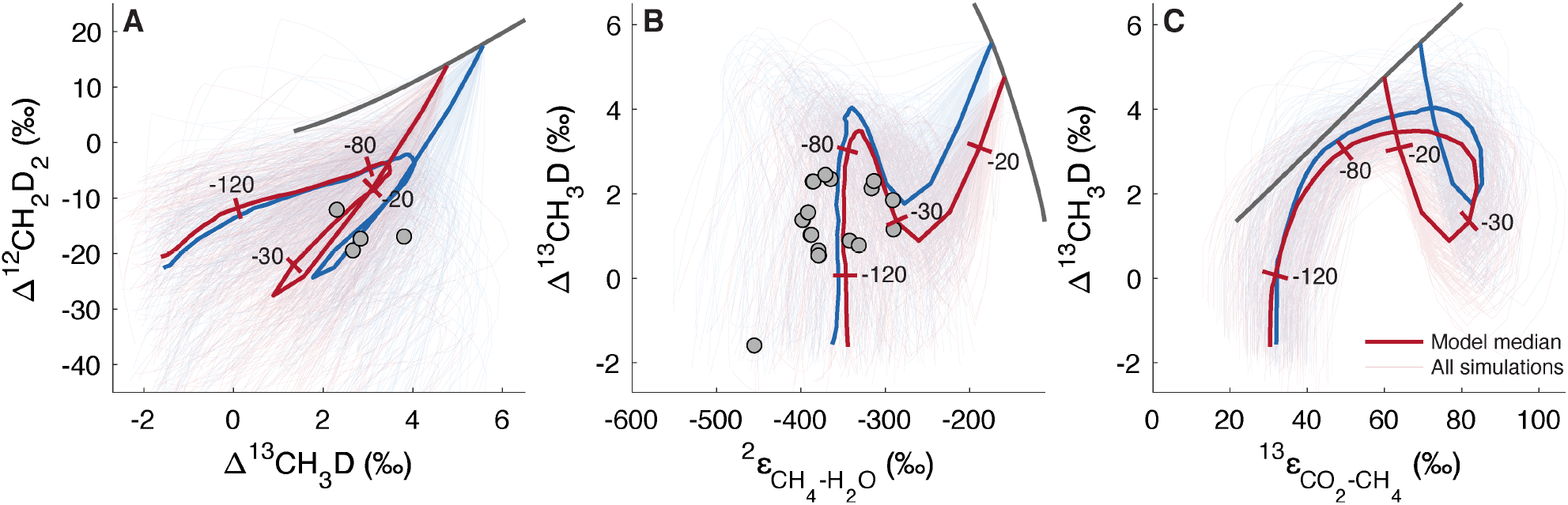
Model-laboratory culture comparison of clumped isotopologue abundances. Experimental data (gray circles) and model results (thin lines) of 200 simulations at 30°C (blue) and 60°C (red). The dark gray lines represent temperature-dependent isotopic equilibrium at 0-350°C, and the thick red and blue lines show the median of the individual simulations, with tickmarks at ΔG_net_ values of –20, –30, –80, and –120 kJmol^−1^. (**A**) Δ^13^CH_3_D against Δ^12^CH_2_D_2_. (**B**) Δ^13^CH_3_D against 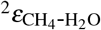. (**C**) Δ^13^CH_3_D against 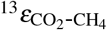. Laboratory culture samples are from hydrogenotrophic methanogens that do not have membrane-associated methanophenazines.

### Isotopic fractionation in energy-limited environments

The metabolic-isotopic model may be used to examine the controls on the isotopic fractionation of methanogenesis not only in laboratory cultures, but in natural environments as well. Laboratory cultures operate far from equilibrium, usually at H_2_ concentrations and cell-specific methanogenesis rates (csMR) much higher than those in natural environments (table S5; fig. S9). Recent reevaluations of slowly forming biogenic methane sources such as marine sediments, coalbeds or shale gas deposits, revealed that apparent CH_4_–CO_2_ and CH_4_–H_2_O isotopic equilibrium is common (8, 25, 37–40). There are currently no laboratory cultures that reproduce the isotopic effects associated with these conditions. We modified the model to use enzyme activities that were measured in H_2_-limited laboratory cultures (Methods) and assessed the resulting ΔG_net_–csMR–isotopic relations at 0-60°C.

Existing estimates of environmental ΔG_net_ are more positive than –30 kJmol^−1^ (table S5). However, the determination of ΔG_net_ in natural environments is often difficult because of low and spatially heterogeneous in-situ H_2_ concentrations (41–43), and the actual range of ΔG_net_ likely reflects this heterogeneity. As ΔG_net_ determines the csMR (fig. S9), which is easier to measure, we henceforth discuss csMR–isotopic fractionation relations. Our model predictions of csMR in energy-limited environments (10^−5^ to 10^0^ fmol cell^−1^ d^−1^) are considerably lower than the typical range in laboratory cultures (10^1^ to 10^4^ fmol cell^−1^ d^−1^, fig. S9, Refs. 23, 29, 44). This csMR difference of several orders of magnitude reflects the model calibration to reproduce H_2_–csMR relations in low-H_2_ laboratory culture experiments (fig. S2). Although this calibration was limited to available experimental H_2_ concentrations higher than 1 *µ*M, our predicted csMR values at ∼nM H_2_ concentrations (10^−5^ to 10^0^ fmol cell^−1^ d^−1^) overlap with the range of csMR values from shallow and deep marine sediments (10^−4^ to 10^0^ fmol cell^−1^ d^−1^), which we calculated from reported bulk methanogenesis rates and cell densities (table S6). We note that there is large uncertainty regarding these environmental csMR values, which stems in part from uncertainty on the net rate of methanogenesis. The methanogenesis rate in natural environments is often determined by radiotracer assays, which may lead to an overestimation of the net methanogenesis rate by orders of magnitude if the reversibility of net methanogenesis (i.e., between CO_2_ and CH_4_) is higher than 0.9 (SM). An additional source of uncertainty is the number of active methanogenic cells in the sediments, and there are currently very limited estimates of these values. Despite these uncertainties, our model predictions of csMR match not only environmental csMR estimates, but also estimates of cell-specific power utilization, *P* = −ΔG_net_ × csMR [W cell^−1^]. We predict *P* between 10^−22^ and 10^−18^ W cell^−1^ for ΔG_net_ between –10 to –30 kJmol^−1^ respectively, at 10°C, in agreement with previously estimated cell-specific power utilization in marine sediments (45).

The model reveals that in contrast to methanogenesis in lab cultures, under energy-limited conditions the Mtr-catalyzed reaction departs from equilibrium before the Mcr-catalyzed reaction (fig. S7). As a consequence, different csMR–fractionation relations emerge, most notably for 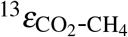 (Fig. 4A). Instead of an increase in 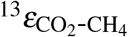 to larger-than-equilibrium values, 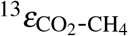 remains approximately constant up to csMR of ∼10 fmol cell^−1^ d^−1^. The reason for this apparent carbon isotopic equilibrium is the similar magnitude of the EFF and KFF of Mtr (17‰ and 16‰ at 60°C, respectively, fig. S4). Thus, as the Mtr-catalyzed reaction departs from equilibrium, the net carbon isotopic fractionation changes little. An environmental prediction of the above is that 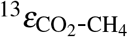 that differs measurably from 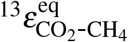 indicates csMR higher than ∼10 fmol cell^−1^ d^−1^.

**Figure 4:**
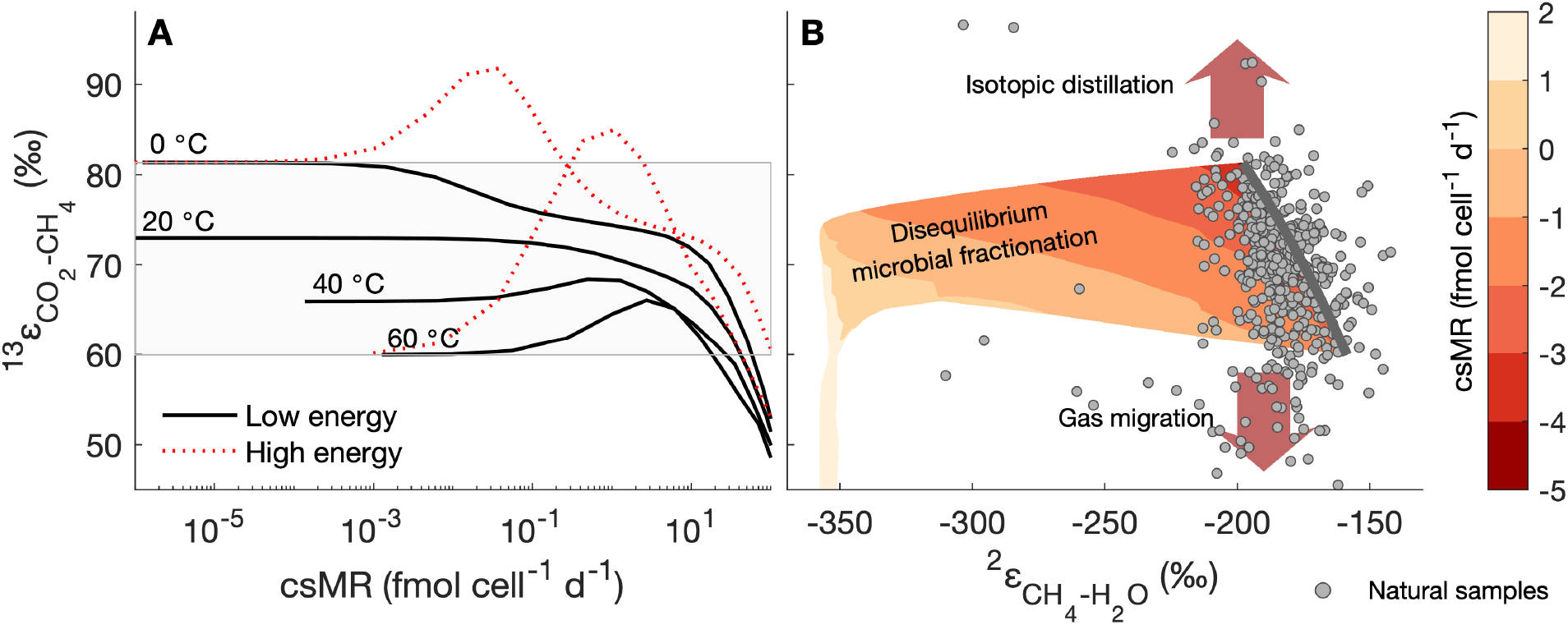
Isotopic fractionation during methanogenesis in energy-limited conditions. (**A**) 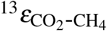 against cell-specific methanogenesis rate (csMR). Dotted red lines show the laboratory-calibrated model (high energy) results for the same temperatures. (**B**) 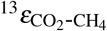 against 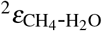. Contours are log_10_(csMR) predicted by our model, between 0°C and 60°C. The calculations are for [H_2_] between 1 nM and 5 *µ*M, [CO_2_] and [CH_4_] of 1 mM, and a cell volume of 1 *µ*m^3^. The circles are biogenic environmental samples from marine sediments, coalbed methane and natural gas deposits (*n* = 491).

Analytically distinguishable departure of 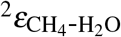 from equilibrium occurs at csMR between ∼0.001 and 0.1 fmol cell^−1^ d^−1^ (at 0°C and 60°C, respectively; fig. S8B), and departure from clumped isotope equilibrium occurs over a similar range of csMR (fig. S8C, D). Thus, our model reveals that at csMR between 0.1 and 10 fmol cell^−1^ d^−1^, one might expect near-equilibrium 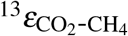 concurrent with disequilibrium 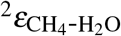. The opposite situation (i.e., near-equilibrium 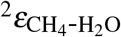 and disequilibrium 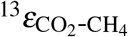) is common in natural environments, though csMR is usually unknown (fig. S9). As suggested in previous studies, this apparent hydrogen isotopic equilibrium concurrent with carbon isotopic disequilibrium may be explained by diffusive mixing of CO_2_ and CH_4_, isotopic (Rayleigh) distillation, or diagenetic isotope exchange without net methane production (Fig. 4B; 8, 38, 46, 47). If disequilibrium 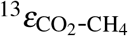 is explained as above rather than by microbial expression of KFFs, then the scarcity of data within the field representing disequilibrium microbial fractionation in Fig. 4B may reflect near-equilibrium isotopic fractionation during methanogenesis in energy-limited environments. This implies csMR lower than ∼0.001– 0.1 fmol cell^−1^ d^−1^(depending on temperature), consistent with almost all csMR estimated from cell abundances and volumetric methanogenesis rates (SM, table S6). In energy-limited environments with higher csMR, near-equilibrium 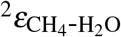 concurrent with disequilibrium 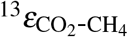 may be explained by methane cycling or anaerobic oxidation of methane that operates close to the thermodynamic limit of this metabolism (48).

## Conclusions

By accounting for the metabolites and reactions in the hydrogenotrophic methanogenesis pathway, we link environmental substrate and product concentrations, pH and temperature to the energetics and net rate of methanogenesis, and to the associated fractionation of carbon, hydrogen and clumped isotopes. The landscape of departure of individual reactions in the pathway from reversibility controls these fractionations, explaining rate–fractionation relations in laboratory cultures. We suggest that a combination of 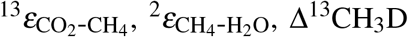 and Δ^12^CH_2_D_2_ data, interpreted within a metabolic-isotopic model framework, can uniquely constrain the in-situ energetics of methanogenic activity (i.e., the actual ΔG_net_ of methanogenesis), which may be difficult to constrain by other means. However, application of such an approach is limited to natural and artificial environments in which methane production dominates the processes that may affect apparent CH_4_–CO_2_ and CH_4_–H_2_O isotopic fractionations. Such processes include CO_2_ and CH_4_ diffusion, isotopic distillation and methane cycling, which are prevalent in natural environments. Nevertheless, the mechanistic understanding of isotopic effects during microbial methanogenesis provided by our model allows an inference that this metabolism operates close to chemical and isotopic equilibrium in a wide range of natural environments, including marine sediments, coal beds and natural gas deposits.

## Methods

### Relating net isotopic fractionation to thermodynamic drive

The metabolic-isotopic model that we developed is based on a general framework, previously used to investigate sulfur isotopes in microbial sulfate-reduction (17, 18). This approach is applicable to any metabolic network with some modifications that we present below. In general, the net flux of a reversible enzymatically catalyzed reaction (*J*) is defined by the difference between the forward (*J*^+^) and backward (*J*^−^) gross fluxes, and can be expressed as (16):

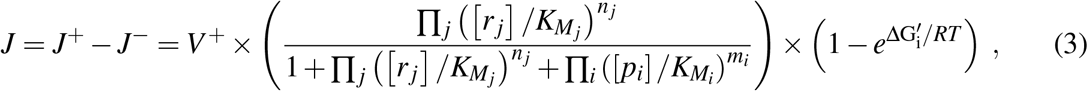

where *V* ^+^ is the maximal rate capacity of the enzyme (defined by 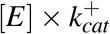, where 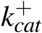 is the catalytic rate constant for forward reaction), [*r* _*j*_] and [*p*_*i*_] are concentrations of the *i*th reactant and *j*th product, respectively, and 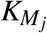 and 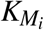 are the Michaelis constants for the forward and reverse directions, respectively. The *n*_*i*_ and *m*_*j*_ stand for the stoichiometry of the reactant and product, respectively. A net flux, as described by Eq. 3, is calculated for each of the reactions in the hydrogenotrophic methanogenesis pathway (table S1). These fluxes are used to construct a mass balance, which is solved at a steady-state to find the concentrations of the metabolites. The steady-state solution was obtained by forward integration using ode15s, an ordinary differential equation solver in MATLAB^®^. As metabolite concentrations span 6 to 7 orders of magnitude, the resulting set of differential equations is stiff, and required the use of this solver with an absolute tolerance of 10^−20^ M. We checked that the duration of integration was sufficient to reach a steady-state solution (i.e., no change in concentrations and fluxes) in all cases.

The model’s inputs include the environmental temperature, pH and extracellular aqueous concentrations of H_2_, CO_2_ and CH_4_. We assume that the intracellular concentrations of H_2_ and CH_4_ are equal to the extracellular concentrations due to their rapid diffusion through the membrane. For CO_2_ we apply a simple diffusion model to relate intracellular to extracellular CO_2_ concentrations. The model’s tunable parameters include enzyme kinetic constants (K_*M*_ and *V* ^+^), thermodynamic constants 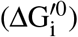 and cellular parameters such as cell size, and concentrations of some of the metabolites. We elaborate on the parameters and the choice of their values in the SM. With the inputs prescribed and values for the tunable parameters chosen, the model outputs are the concentrations of all intracellular metabolites and the gross fluxes among these metabolites, which are related to the reactions in the methanogenesis pathway.

We used the forward and backward gross fluxes from the metabolic model to calculate the net isotopic fractionations in hydrogenotrophic methanogenesis. To this end, we constructed a mass balance for each isotopic system (for both bulk and clumped isotopes), which is based on calculating an isotopic flux associated with each of the chemical fluxes. For the schematic reaction *r* → *p*, this isotopic flux can be approximated:

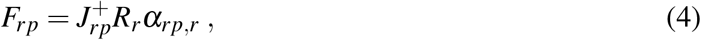

where 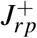 is the forward chemical flux between metabolites pools *r* and *p, R*_*r*_ is the abundance ratio of heavy to light isotopes in pool *r*, and *α*_*rp,r*_ is the KFF of the reaction. A term similar to Eq. 4 was assigned to account for the consumption and production of each isotopologue in the pathway while considering the stoichiometry and the symmetry coefficients, where relevant, in a similar manner to previous model derivations (7, 9). As described above, these terms were used to construct isotopic mass balances in the form of a set of coupled differential equations, which were solved numerically using the ode15s solver in MATLAB^®^ to obtain the steady-state bulk and clumped isotopic composition of all intracellular metabolites, given bulk extracellular D/H and ^13^C/^12^C of H_2_O and CO_2_, respectively.

The net isotopic fractionation in linear metabolic networks such as the carbon reaction network in hydrogenotrophic methanogenesis can be solved analytically. For example, in the case of the reaction *r* += *p*, the net isotopic fractionation between the metabolite pools of *r* and 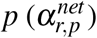 can be calculated by:

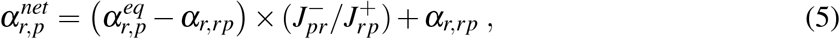

where 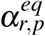 is the EFF between pools *r* and *p, α* _*r,rp*_ is 1/KFF of the forward reaction. A full derivation for this term is presented in Wing and Halevy, 2014 (18). The reversibility 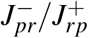 is directly related to 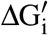 through the flux-force relationship (Eq. 2). In the case of longer linear reaction networks such as *s* ⇆ *r* ⇆ *p*, the net isotopic fractionation can be described by:

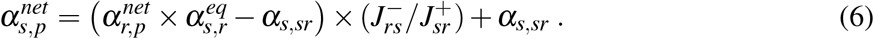

This expression can be expanded recursively to describe the net isotopic fractionation in a linear reaction network of arbitrary length (18). Such a recursive expression has the advantage of a significantly reduced computational time compared to the numerical solution that is used for nonlinear reaction networks. We verified that this analytical solution yields identical results to the numerical solution for the net carbon isotopic fractionation.

### Isotopic model parameters

To calculate the net isotopic fractionations associated with each of the steps in the methanogenesis pathway we used the temperature-dependent EFFs which were calculated for the hydrogenotrophic pathway (25, 49). The KFFs were determined experimentally only for Mcr (24). To account for missing KFFs we randomly sampled their values from prior uniform distributions 10^6^ times. We included both primary and secondary isotopic effects (*α*_*p*_ and *α*_*s*_, respectively). Primary isotopic effects are due to the breaking or formation of a bond directly with the atom of interest, and secondary isotopic effects are due to breaking or formation of an adjacent bond. Secondary KFFs are usually smaller than primary KFFs. We weighted the combinations of KFFs drawn from the prior distributions by the model-experimental mismatch, expressed as the inverse of the square of sum of the square errors (1/SSE^2^), to generate posterior distributions of the KFFs, which were then used in the model. The selection of prior KFF value distributions is described in the SM.

The posterior distributions of the KFFs (fig. S4) serve as a sensitivity analysis of the model. Posterior distributions that are similar to the prior distributions indicate insensitivity to the model parameter, in this case, the KFF value. For carbon isotopes, the model shows sensitivity to the KFFs of Fmd, Ftr and Mtr with smallest-mismatch values of 30.8‰, 25.8‰, and 15.8‰ at mesophilic conditions, respectively. This sensitivity is in line with our observations of the reactions that determine the trajectory of 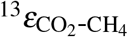 departure from equilibrium. For hydrogen isotopes, the model shows sensitivity to most of the primary KFFs, and to the secondary KFF of Mtr. Notably, the primary KFFs of Mtd, Mer and Frh tend to be small (i.e., closer to unity), with median values between ≈–450‰ and ≈–300‰, but larger than the secondary KFFs with median values between ≈–330‰ and ≈–220‰ .

### Simulating methanogenesis in energy-limited conditions

The metabolic parameters that we used to explore isotopic fractionation in methanogenesis were curated from cell cultures grown under optimal conditions. However, methanogens tightly regulate gene expression under energy-limiting conditions, resulting in shifts in enzyme specific activities (30, 50–57). Specifically, prolonged H_2_ limitation promotes higher activities of Frh, Mvh/Hdr and the Mcr I isoenzyme, in parallel to a decrease in the activity of Mcr II (52, 53, 58). Mcr I has a higher affinity to the substrates CH_3_-SCoM and HS-CoB relative to Mcr II, allowing the cells to increase the cell-specific respiration rate when H_2_ becomes limiting (59–61). Therefore, in the simulations of methanogenesis in natural, energy-limited environments we used enzyme activities measured in crude extracts from cells that were grown under conditions favoring Mcr I activity (53) with *K*_*M*_ values of Mcr I (table S2). In laboratory cultures, cells under energy limitation decrease their volume in a matter of days or weeks, and a similar phenomenon was observed when comparing cell communities in energy-replete *vs*. energy-deprived environmental settings (62). Thus, we use a cellular volume smaller by a factor of two from the default value (2 fL cell^−1^ *vs*. 1 fL cell^−1^).

## Acknowledgments

J.G. acknowledges support from the Sustainability and Energy Research Initiative (SAERI) of the Weizmann Institute of Science. Q.J. is supported by NSF award EAR-1636815.

## Author contributions

J.G. and I.H. conceived the study, developed and analyzed the metabolic-isotopic model, and wrote the initial draft. Q.J. developed the metabolic model. All authors contributed to the writing.

## References

[1] M. Saunois, et al., The global methane budget 2000–2012. Earth Syst. Sci. Data 8, 697–751 (2016).

[2] S. Schwietzke, O. A. Sherwood, L. M. P. Bruhwiler, J. B. Miller, G. Etiope, E. J. Dlugokencky, S. E. Michel, V. A. Arling, B. H. Vaughn, J. W. C. White, P. P. Tans, Upward revision of global fossil fuel methane emissions based on isotope database. Nature 538, 88–91 (2016).

[3] Z. Lyu, N. Shao, T. Akinyemi, W. B. Whitman, Methanogenesis. Current Biology 28, R727–R732 (2018).

[4] M. J. Whiticar, Carbon and hydrogen isotope systematics of bacterial formation and oxidation of methane. Chem. Geol. 161, 291–314 (1999).

[5] E. G. Nisbet, E. J. Dlugokencky, M. R. Manning, D. Lowry, R. E. Fisher, J. L. France, S. E. Michel, J. B. Miller, J. W. C. White, B. Vaughn, P. Bousquet, J. A. Pyle, N. J. Warwick, M. Cain, R. Brownlow, G. Zazzeri, M. Lanoisellé, A. C. Manning, E. Gloor, D. E. J. Worthy, E.-G. Brunke, C. Labuschagne, E. W. Wolff, A. L. Ganesan, Rising atmospheric methane: 2007-2014 growth and isotopic shift. Glob. Biogeochem. Cycles 30, 1356–1370 (2016).

[6] D. A. Stolper, A. Sessions, A. Ferreira, E. Santos Neto, A. Schimmelmann, S. Shusta, D. Valentine, J. Eiler, Combined 13C–D and D–D clumping in methane: Methods and preliminary results. Geochim. Cosmochim. Acta 126, 169–191 (2014).

[7] D. T. Wang, D. S. Gruen, B. S. Lollar, K.-U. Hinrichs, L. C. Stewart, J. F. Holden, A. N. Hristov, J. W. Pohlman, P. L. Morrill, M. Könneke, K. B. Delwiche, E. P. Reeves, C. N. Sutcliffe, D. J. Ritter, J. S. Seewald, J. C. McIntosh, H. F. Hemond, M. D. Kubo, D. Cardace, T. M. Hoehler, S. Ono, Nonequilibrium clumped isotope signals in microbial methane. Science 348, 428–431 (2015).

[8] T. Okumura, S. Kawagucci, Y. Saito, Y. Matsui, K. Takai, H. Imachi, Hydrogen and carbon isotope systematics in hydrogenotrophic methanogenesis under H2-limited and H2-enriched conditions: Implications for the origin of methane and its isotopic diagnosis. Prog. Earth Planet. Sci. 3, 14 (2016).

[9] X. Cao, H. Bao, Y. Peng, A kinetic model for isotopologue signatures of methane generated by biotic and abiotic CO2 methanation. Geochim. Cosmochim. Acta 249, 59–75 (2019).

[10] T. Wagner, U. Ermler, S. Shima, The methanogenic CO2 reducing-and-fixing enzyme is bifunctional and contains 46 [4Fe-4S] clusters. Science 354, 114–117 (2016).

[11] A.-K. Kaster, J. Moll, K. Parey, R. K. Thauer, Coupling of ferredoxin and heterodisulfide reduction via electron bifurcation in hydrogenotrophic methanogenic archaea. PNAS 108, 2981–6 (2011).

[12] L. M. I. de Poorter, W. J. Geerts, J. T. Keltjens, Hydrogen concentrations in methane-forming cells probed by the ratios of reduced and oxidized coenzyme F420. Microbiology 151, 1697–1705 (2005).

[13] L. M. I. de Poorter, W. G. Geerts, A. P. R. Theuvenet, J. T. Keltjens, Bioenergetics of the formyl-methanofuran dehydrogenase and heterodisulfide reductase reactions in Methanothermobacter thermautotrophicus. Eur. J. Biochem. 270, 66–75 (2002).

[14] J. Bassham, G. Krause, Free energy changes and metabolic regulation in steady-state photosynthetic carbon reduction. Biochim. Biophys. Acta BBA - Bioenerg. 189, 207–221 (1969).

[15] D. A. Beard, H. Qian, Relationship between thermodynamic driving force and one-way fluxes in reversible processes. PloS One 2, e144 (2007).

[16] E. Noor, A. Flamholz, W. Liebermeister, A. Bar-Even, R. Milo, A note on the kinetics of enzyme action: A decomposition that highlights thermodynamic effects. FEBS Lett. 587, 2772–7 (2013).

[17] C. Rees, A steady-state model for sulphur isotope fractionation in bacterial reduction processes. Geochim. Cosmochim. Acta 37, 1141–1162 (1973).

[18] B. A. Wing, I. Halevy, Intracellular metabolite levels shape sulfur isotope fractionation during microbial sulfate respiration. Proc. Natl. Acad. Sci. U. S. A. 111, 18116–25 (2014).

[19] D. L. Valentine, A. Chidthaisong, A. Rice, W. S. Reeburgh, S. C. Tyler, Carbon and hydrogen isotope fractionation by moderately thermophilic methanogens. Geochim. Cosmochim. Acta 68, 1571–1590 (2004).

[20] H. Penning, C. M. Plugge, P. E. Galand, R. Conrad, Variation of carbon isotope fractionation in hydrogenotrophic methanogenic microbial cultures and environmental samples at different energy status. Glob. Change Biol. 11, 2103–2113 (2005).

[21] K. Takai, K. Nakamura, T. Toki, U. Tsunogai, M. Miyazaki, J. Miyazaki, H. Hirayama, S. Nakagawa, T. Nunoura, K. Horikoshi, Cell proliferation at 122 degrees C and isotopically heavy CH4 production by a hyperthermophilic methanogen under high-pressure cultivation. Proc. Natl. Acad. Sci. U. S. A. 105, 10949–54 (2008).

[22] S. Hattori, H. Nashimoto, H. Kimura, K. Koba, K. Yamada, M. Shimizu, H. Watanabe, M. Yoh, N. Yoshida, Hydrogen and carbon isotope fractionation by thermophilic hydrogenotrophic methanogens from a deep aquifer under coculture with fermenters. Geochem. J. 46, 193–200 (2012).

[23] B. D. Topçuoğlu, C. Meydan, T. B. Nguyen, S. Q. Lang, J. F. Holden, Growth Kinetics, Carbon Isotope Fractionation, and Gene Expression in the Hyperthermophile Methanocaldococcus jannaschii during Hydrogen-Limited Growth and Interspecies Hydrogen Transfer. Appl. Environ. Microbiol. 85, 1–14 (2019).

[24] S. Scheller, M. Goenrich, R. K. Thauer, B. Jaun, Methyl-coenzyme M reductase from methanogenic archaea: Isotope effects on the formation and anaerobic oxidation of methane. J. Am. Chem. Soc. 135, 14975–84 (2013).

[25] J. Gropp, M. A. Iron, I. Halevy, Theoretical estimates of equilibrium carbon and hydrogen isotope effects in microbial methane production and anaerobic oxidation of methane. Geochimica et Cosmochimica Acta 295, 237–264 (2021).

[26] H. M. Miller, N. Chaudhry, M. E. Conrad, M. Bill, S. H. Kopf, A. S. Templeton, Large carbon isotope variability during methanogenesis under alkaline conditions. Geochim. Cosmochim. Acta 237, 18–31 (2018).

[27] D. B. Nothaft, A. S. Templeton, J. H. Rhim, D. T. Wang, J. Labidi, H. M. Miller, E. S. Boyd, J. M. Matter, S. Ono, E. D. Young, S. H. Kopf, P. B. Kelemen, M. E. Conrad, Geochemical, biological and clumped isotopologue evidence for substantial microbial methane production under carbon limitation in serpentinites of the Samail Ophiolite, Oman. J. Geophys. Res. Biogeosciences NA, e2020JG006025 (2021).

[28] H. Yoshioka, S. Sakata, Y. Kamagata, Hydrogen isotope fractionation by Methanothermobacter thermoautotrophicus in coculture and pure culture conditions. Geochim. Cosmochim. Acta 72, 2687–2694 (2008).

[29] S. Kawagucci, M. Kobayashi, S. Hattori, K. Yamada, Y. Ueno, K. Takai, N. Yoshida, Hydrogen isotope systematics among H2–H2O–CH4 during the growth of the hydrogenotrophic methanogen Methanothermobacter thermautotrophicus strain ΔH. Geochim. Cosmochim. Acta 142, 601–614 (2014).

[30] C. Afting, E. Kremmer, C. Brucker, A. Hochheimer, R. K. Thauer, Regulation of the synthesis of H2-forming methylenetetrahydromethanopterin dehydrogenase (Hmd) and of HmdII and HmdIII in Methanothermobacter marburgensis. Arch. Microbiol. 174, 225–232 (2000).

[31] R. Botz, H. D. Pokojski, M. Schmitt, M. Thomm, Carbon isotope fractionation during bacterial methanogenesis by CO2 reduction. Org. Geochem. 25, 255–262 (1996).

[32] P. M. Douglas, D. A. Stolper, J. M. Eiler, A. L. Sessions, M. Lawson, Y. Shuai, A. Bishop, O. G. Podlaha, A. A. Ferreira, E. V. Santos Neto, M. Niemann, A. S. Steen, L. Huang, L. Chimiak, D. L. Valentine, J. Fiebig, A. J. Luhmann, W. E. Seyfried, G. Etiope, M. Schoell, W. P. Inskeep, J. J. Moran, N. Kitchen, Methane clumped isotopes: Progress and potential for a new isotopic tracer. Org. Geochem. 113, 262–282 (2017).

[33] D. A. Stolper, A. Martini, M. Clog, P. Douglas, S. Shusta, D. Valentine, A. Sessions, J. Eiler, Distinguishing and understanding thermogenic and biogenic sources of methane using multiply substituted isotopologues. Geochim. Cosmochim. Acta 161, 219–247 (2015).

[34] L. Y. Yeung, Combinatorial effects on clumped isotopes and their significance in biogeochemistry. Geochim. Cosmochim. Acta 172, 22–38 (2016).

[35] T. Röckmann, M. E. Popa, M. C. Krol, M. E. G. Hofmann, Statistical clumped isotope signatures. Sci. Rep. 6, 31947 (2016).

[36] L. Taenzer, J. Labidi, A. L. Masterson, X. Feng, D. Rumble, E. D. Young, W. D. Leavitt, Low Δ12CH2D2 values in microbialgenic methane result from combinatorial isotope effects. Geochimica et Cosmochimica Acta 285, 225–236 (2020).

[37] D. S. Vinson, N. E. Blair, A. M. Martini, S. Larter, W. H. Orem, J. C. McIntosh, Microbial methane from in situ biodegradation of coal and shale: A review and reevaluation of hydrogen and carbon isotope signatures. Chem. Geol. 453, 128–145 (2017).

[38] A. C. Turner, R. Korol, D. L. Eldridge, M. Bill, M. E. Conrad, T. F. Miller, D. A. Stolper, Experimental and theoretical determinations of hydrogen isotopic equilibrium in the system CH4-H2-H2O from 3 to 200°C. Geochimica et Cosmochimica Acta NA, NA (2021).

[39] J. J. Jautzy, P. M. J. Douglas, H. Xie, J. M. Eiler, I. D. Clark, CH4 isotopic ordering records ultra-slow hydrocarbon biodegradation in the deep subsurface. Earth and Planetary Science Letters 562, 116841 (2021).

[40] N. Zhang, G. T. Snyder, M. Lin, M. Nakagawa, A. Gilbert, N. Yoshida, R. Matsumoto, Y. Sekine, Doubly substituted isotopologues of methane hydrate (13CH3D and 12CH2D2): Implications for methane clumped isotope effects, source apportionments and global hydrate reservoirs. Geochimica et Cosmochimica Acta NA, NA (2021).

[41] R. Conrad, T. J. Phelps, J. G. Zeikus, Gas metabolism evidence in support of the juxtaposition of hydrogen-producing and methanogenic bacteria in sewage sludge and lake sediments. Appl. Environ. Microbiol. 50, 595–601 (1985).

[42] E. Giraldo-Gomez, S. Goodwin, M. S. Switzenbaum, Influence of mass transfer limitations on determination of the half saturation constant for hydrogen uptake in a mixed-culture CH4producing enrichment. Biotechnol. Bioeng. 40, 768–76 (1992).

[43] Y.-S. Lin, V. B. Heuer, T. Goldhammer, M. Y. Kellermann, M. Zabel, K.-U. Hinrichs, Towards constraining H2 concentration in subseafloor sediment: A proposal for combined analysis by two distinct approaches. Geochimica et Cosmochimica Acta 77, 186–201 (2012).

[44] A. M. Zyakun, Potential of 13C/12C Variations in Bacterial Methane in Assessing Origin of Environmental Methane. Hydrocarbon Migration And Its Near-Surface Expression, D. Schumacher, M. A. Abrams, eds. (AAPG Special Volumes, 1996), pp. 341–352.

[45] J. A. Bradley, S. Arndt, J. P. Amend, E. Burwicz, A. W. Dale, M. Egger, D. E. LaRowe, Widespread energy limitation to life in global subseafloor sediments. Sci. Adv. 6, eaba0697 (2020).

[46] C. K. Paull, T. D. Lorenson, W. S. Borowski, W. Ussler III, K. Olsen, N. M. Rodriguez, Isotopic Composition of CH4, CO2 Species, and Sedimentary Organic Matter Within Samples From the Blake Ridge: Gas Source Implications. Proc. Ocean Drill. Program Sci. Results 164, 67–78 (2000).

[47] J. W. Pohlman, M. Kaneko, V. B. Heuer, R. B. Coffin, M. Whiticar, Methane sources and production in the northern Cascadia margin gas hydrate system. Earth and Planetary Science Letters 287, 504–512 (2009).

[48] G. Wegener, J. Gropp, H. Taubner, I. Halevy, M. Elvert, Sulfate-dependent reversibility of intracellular reactions explains the opposing isotope effects in the anaerobic oxidation of methane. Sci. Adv. 7, eabe4939 (2021).

[49] J. Gropp, M. A. Iron, I. Halevy, Corrigendum to “Theoretical estimates of equilibrium carbon and hydrogen isotope effects in microbial methane production and anaerobic oxidation of methane” [Geochim. Cosmochim. Acta 295 (2021) 237–264]. Geochimica et Cosmochimica Acta 306, 386–389 (2021).

[50] R. M. Morgan, T. D. Pihl, J. Nölling, J. N. Reeve, Hydrogen regulation of growth, growth yields, and methane gene transcription in Methanobacterium thermoautotrophicum deltaH. J. Bacteriol. 179, 889–98 (1997).

[51] J. N. Reeve, J. Nölling, R. M. Morgan, D. R. Smith, Methanogenesis: Genes, genomes, and who’s on first? J. Bacteriol. 179, 5975–86 (1997).

[52] P. Vermeij, J. L. Pennings, S. M. Maassen, J. T. Keltjens, G. D. Vogels, Cellular levels of factor 390 and methanogenic enzymes during growth of Methanobacterium thermoautotrophicum deltaH. J. Bacteriol. 179, 6640–8 (1997).

[53] J. L. Pennings, P. Vermeij, L. M. de Poorter, J. T. Keltjens, G. D. Vogels, Adaptation of methane formation and enzyme contents during growth of Methanobacterium thermoautotrophicum (strain ΔH) in a fed-batch fermentor. Antonie Van Leeuwenhoek 77, 281–291 (2000).

[54] E. L. Hendrickson, A. K. Haydock, B. C. Moore, W. B. Whitman, J. A. Leigh, Functionally distinct genes regulated by hydrogen limitation and growth rate in methanogenic Archaea. Proc. Natl. Acad. Sci. U. S. A. 104, 8930–4 (2007).

[55] S. Kato, T. Kosaka, K. Watanabe, Comparative transcriptome analysis of responses of Methanothermobacter thermautotrophicus to different environmental stimuli. Environ. Microbiol. 10, 893–905 (2008).

[56] Q. Xia, T. Wang, E. L. Hendrickson, T. J. Lie, M. Hackett, J. A. Leigh, Quantitative proteomics of nutrient limitation in the hydrogenotrophic methanogen Methanococcus maripaludis. BMC Microbiol. 9, 149 (2009).

[57] M. Enoki, N. Shinzato, H. Sato, K. Nakamura, Y. Kamagata, Comparative Proteomic Analysis of Methanothermobacter themautotrophicus ΔH in Pure Culture and in Co-Culture with a Butyrate-Oxidizing Bacterium. PLoS ONE 6, e24309 (2011).

[58] T. D. Pihl, S. Sharma, J. N. Reeve, Growth phase-dependent transcription of the genes that encode the two methyl coenzyme M reductase isoenzymes and N5-methyltetrahydromethanopterin:coenzyme M methyltransferase in Methanobacterium thermoautotrophicum delta H. J. Bacteriol. 176, 6384–6391 (1994).

[59] L. G. Bonacker, S. Baudner, R. K. Thauer, Differential expression of the two methylcoenzyme M reductases in Methanobacterium thermoautotrophicum as determined immunochemically via isoenzyme-specific antisera. Eur. J. Biochem. 206, 87–92 (1992).

[60] L. G. Bonacker, S. Baudner, E. Mörschel, R. Böcher, R. K. Thauer, Properties of the two isoenzymes of methyl-coenzyme M reductase in Methanobacterium thermoautotrophicum. Eur. J. Biochem. 217, 587–95 (1993).

[61] M. Dey, X. Li, R. C. Kunz, S. W. Ragsdale, Detection of Organometallic and Radical Intermediates in the Catalytic Mechanism of Methyl-Coenzyme M Reductase Using the Natural Substrate Methyl-Coenzyme M and a Coenzyme B Substrate Analogue. Biochemistry 49, 10902–10911 (2010).

[62] M. A. Lever, K. L. Rogers, K. G. Lloyd, J. Overmann, B. Schink, R. K. Thauer, T. M. Hoehler, B. B. Jørgensen, Life under extreme energy limitation: A synthesis of laboratory-and field-based investigations. FEMS Microbiol Rev 39, 688–728 (2015).

