## Supplementary material for "Controls on the isotopic composition of microbial methane": SM

### Supplementary Materials

#### Metabolic model parameters

We tested the model for input $H_2$(aq) concentration ranges of $10^{-9}$ to $10^{-2}$ M, $CO_2$(aq) concentration ranges of 1 to 100 mM, and $CH_4$(aq) was held at $10^{-5}$ M. These ranges were chosen to include concentrations reported in various culture experiments, and guided by a sensitivity analysis, which showed that the model is relatively insensitive to $CH_4$(aq) levels but is sensitive to $H_2$(aq) and $CO_2$(aq) levels. We numerically solve for the steady-state concentrations of 16 metabolites in the hydrogenotrophic methanogenesis pathway of *Methanothermobacter thermoautotrophicus*. The initial concentrations of the total (i.e., reduced and oxidized) electron carriers (ferredoxin, coenzyme B, $F_{420}$) and $H_4$MPT were compiled from experimentally determined concentrations, where available (table S3).

##### Extracellular and intracellular concentrations of $CO_2$, $H_2$ and $CH_4$

The model was designed to predict intracellular metabolite concentrations of the hydrogenotrophic methanogenesis pathway at a steady state, by prescribing temperature, and extracellular concentrations of $H_2$(aq), $CO_2$(aq) and $CH_4$(aq). These small, non-polar molecules diffuse rapidly through the cell membrane, and it is often assumed that their intracellular aqueous concentrations are in equilibrium with the headspace gas (related to the aqueous concentrations through Henry's Law constants; table S3). We adopt this assumption for $CH_4$, to which the model is relatively insensitive. However, we explore two cases in which concentrations of $CO_2$ and $H_2$ are not in equilibrium with the headspace:

(*i*) $CO_2$ diffusion through the membrane may become limiting with increasing net methanogenesis flux (1). We model intracellular $CO_2$ concentrations $[CO_{2(in)}]$ using Fick's law:

$$J_{dif} = D \times A \times \frac{[CO_{2(out)}] - [CO_{2(in)}]}{z}, \qquad \text{(S1)}$$

where $J_{dif}$ is the net $CO_2$ diffusive flux, $D$ is the permeability constant, $A$ is the membrane surface area, $z$ is the membrane width and $[CO_{2(out)}]$ is the extracellular $CO_2$ concentration.

(*ii*) Cultivation of methanogens under low $H_2$(aq) conditions requires growth in co-cultures with $H_2$-producing (hydrogenic) bacteria (2, 3). Juxtaposition of methanogens and hydrogenic bacteria may result in immediate consumption of bacterially produced $H_2$ by the methanogens, which may result in spatially heterogeneous $H_2$ concentrations (4, 5). In this case, the aqueous $H_2$ concentration in the immediate vicinity of the methanogens may be higher than the $H_2$ concentration in equilibrium with the headspace. The best fit between the net carbon isotopic fractionation in methanogenesis ($^{13}\varepsilon_{CO_2\text{-}CH_4}$) measured in laboratory co-cultures and predicted in our model is obtained at in-situ $H_2$ concentrations that are approximately five times higher than the concentration at equilibrium with the headspace.

#### Energy conservation

Methanogens conserve chemical energy by coupling the exergonic methyl-group transfer by the Mtr-catalyzed reaction to export of $Na^+$ ions out of the cell. This generates an electrochemical gradient, which is used to phosphorylate ADP to ATP by a proton- or sodium-driven ATPase (6, 7). Hydrogenotrophic organisms without membrane-associated methanophenazines such as *M. thermoautotrophicus*, have an ATP yield ($Y_{ATP}$) of $\approx$0.5 mole ATP per mole $CH_4$ under optimal growth conditions (8). Production of 1 mole ATP in physiological conditions requires +43.5 kJ (9), and if we assume that the energy required for its production comes solely from $Na^+$ translocation by Mtr, 0.5 moles of ATP requires $\approx$22 kJ. Based on the observations that the minimal threshold for methanogenic activity is –10 $kJ\,mol^{-1}$, we estimate that under non-optimal growth conditions $Y_{ATP}$ is closer to 0.2, requiring at least 9 $kJ\,mol^{-1}$. We thus chose a constant $Y_{ATP}$ of 0.2.

#### Enzyme kinetic parameters

The Michaelis constants ($K_M$) were compiled from the published literature (table S2). We explore the sensitivity to these values in figures S11-S13. Some $K_M$ values were missing in the literature. Based on the observation that $K_M$ values in various organisms are normally distributed (10), we randomly sampled ten missing $K_M$ values over $10^5$ simulations from prescribed prior normal distributions. Wherever possible, we determined the mean and $1\sigma$ values of these prior distributions based on metabolite-specific $K_M$ values from other enzymes, deposited in the BRENDA database. We then generated posterior distributions for each $K_M$ value, by considering the inverse of the sum of squared errors ($\mathrm{SSE}^{-1}$) between the model results and the experimental measurements of biomass-specific methanogenesis rate and their dependence on $H_2$ concentrations (11) (fig. S14). We use the posterior distributions to generate the uncertainty envelopes of the departure from equilibrium (Fig. 1B). For the isotopic model we use the median $K_M$ values from each posterior distribution.

The maximal rate capacity $V^+$ that is used in Eq. 3 is a product of enzyme concentrations ($[E]$) and turnover rates ($k^+_{cat}$), and even if all $k^+_{cat}$ values are known, calculation of $V^+$ is not possible without knowledge of enzyme levels under various physiological conditions. Instead, we chose to use specific activities determined from pure culture crude cell extracts ($V^+_{vitro}$), which provide an approximation of $k^+_{cat} \times [E]$ of all enzymes in the pathway. We explore the sensitivity to these values in figures S11-S13. Although this approach avoids the need to prescribe total enzyme concentrations and to know $k^+_{cat}$ of all of the enzymes in the pathway, $V^+_{vitro}$ does not necessarily reflect in-vivo activities ($V^+_{vivo}$). To account for the difference between in-vivo and in-vitro conditions we used a scaling factor:

$$U_{viv/vit} = \frac{[E]_{vivo} \times k^+_{cat,vivo}}{[E]_{vitro} \times k^+_{cat,vitro}}. \qquad \text{(S2)}$$

Assuming that $k^+_{cat,vitro} \approx k^+_{cat,vivo}$, this term reduces to $U_{viv/vit} = [E]_{vivo}/[E]_{vitro}$, which implies that this scaling factor is essentially a ratio of the active enzyme concentrations of in-vitro and in-vivo conditions. We also assumed that this ratio is uniform across all enzymes. To determine $U_{viv/vit}$ we used the model-experiment mismatch as was described for the $K_M$ values (fig. S14). We found that using a $U_{viv/vit}$ value of $\sim$6 yields the best match between the model and experimental results of biomass-specific methanogenesis rates.

#### Temperature effects on enzyme kinetics

In our model we use enzyme activities taken from crude cell extracts that were mostly cultured at 60°C. The maximal rate capacity of an enzyme at a certain temperature ($V^+_2$) is a function of its activity at the reference temperature ($V_1^+$), the difference between the reference temperature and the temperature of interest ($\Delta$T), and the scaling parameter $Q_{10}^{V^+}$:

$$V_2^+ = V_1^+ \times \left(Q_{10}^{V^+}\right)^{\Delta T/10°C} . \tag{S3}$$

The overall methanogenic rate is expected to increase with temperature, and this dependency is often described by the empirical temperature coefficient $Q_{10}$, which reflects the effect of temperature on both the enzyme kinetics ($Q_{10}^{V^+}$) and the thermodynamic drive $\Delta G_{net}$. Previous estimates of $Q_{10}$ range between 1 and 10, depending on environmental parameters, such as the temperature and substrate availability (12, 13). To calculate $Q_{10}$ we use the ratio of the cell-specific rates of methanogenesis from the model ($J_i$) at a specific $\Delta G_{net}$ value: $Q_{10} = (J_2/J_1)^{(10°C/\Delta T)}$ (fig. S9). We found that for $\Delta G_{net}$ values between –10 and –60 kJ mol$^{-1}$, $Q_{10}$ is between $\sim$3 for $Q_{10}^{V^+}$= 1, and $\sim$6 for $Q_{10}^{V^+}$ = 3 (fig. S10). Both these values are in agreement with the ranges observed in natural environments (12). We explored how the choice of $Q_{10}^{V^+}$ affects the $^{13}\varepsilon_{CO_2\text{-}CH_4}$ relation to the cell-specific rates of methanogenesis (csMR) by comparing our model simulations to experiments from Botz et al., 1996 (14). We found that a $Q_{10}^{V^+}$ values of 1 yields an optimal fit, and that the main parameter that controls the $^{13}\varepsilon_{CO_2\text{-}CH_4}$–csMR mismatch was the cell size rather than $Q_{10}^{V^+}$ (S17). We therefore chose to use a $Q_{10}^{V^+}$ value of 1 in the model.

#### Isoenzymes of methylene-$H_4$MPT dehydrogenase

The methenyl-$H_4$MPT reduction step in the hydrogenotrophic methanogenesis pathway can be catalyzed by either $F_{420}H_2$- or $H_2$-dependent hydrogenases (Mtd or Hmd, respectively). There is comprehensive evidence that these enzymes are expressed in a mutually exclusive manner (15–17). In certain conditions, such as exponential growth in $H_2$-replete media, Hmd is up to 25 times more active than Mtd, while in $H_2$-limiting conditions Mtd is up to 25 times more active (15). Hmd catalyzes the reduction of methylene-$H_4$MPT by a hydride ion directly from $H_2$, and at high enough activities it may introduce a mixing effect on the net $^2\varepsilon_{CH_4\text{-}H_2O}$ and a combinatorial effect on $\Delta^{12}CH_2D_2$. To explore these mixing effects we switched the Hmd activities in the default model to activities that were measured under conditions that promote high expression of Hmd.

#### Isotopic model parameters

We used temperature-dependent equilibrium fractionation factors (EFFs) that were calculated for the hydrogenotrophic pathway (18, 19), and experimentally-measured kinetic fractionation factors (KFFs) where available (20). To account for missing KFFs we randomly sampled their values from prior uniform distributions over $10^6$ simulations. We included both primary and secondary isotopic effects ($\alpha_p$ and $\alpha_s$, respectively). We assumed normal KFFs (i.e., the light isotopologues react faster than the heavy isotopologues, $\alpha < 1$) for both carbon and hydrogen isotopes, and assigned $\alpha_p \in (0.2, 1)$ and $\alpha_s \in (0.6, 1)$ for hydrogen isotopes, and $\alpha_p \in (0.95, 1)$ and $\alpha_s \in (0.95, 1)$ for carbon isotopes. Notably, optimal model-measurement fits were obtained with an inverse KFF (i.e., $\alpha > 1$) for the Mvh/Hdr-catalyzed back reaction. While normal KFFs are more common, in some cases inverse KFFs emerge, often in a multistep enzymatic reaction that is treated as one composite reaction. In such reactions, chemical and isotopic equilibrium between reaction intermediates before the rate determining steps may lead to an overall (apparent) inverse KFF, even if the rate determining step itself has a normal KFF (21–23).

The KFFs of Mcr were assigned distributions based on the experimental measurements. The carbon KFF from $CH_3$-SCoM to $CH_4$ was drawn from a uniform distribution $\alpha \in (0.952, 0.97)$, and the hydrogen KFFs were drawn from normal distributions (for $\alpha_p$ $\mu = 0.41$, $\sigma = 0.04$ and for $\alpha_s$ $\mu = 0.85$, $\sigma = 0.035$). We weighted the combinations of KFFs drawn from these prior distributions by the model-experimental mismatch to the sum of squared errors SSE (using $1/\text{SSE}^2$) to generate posterior distributions of the KFFs, which were then used in the model (fig. S4).

Under the product rule, the clumped isotopologue KFFs are the product of the individual KFFs of the two reactions that generate the clumped isotopologue. However, there is evidence that in some cases these KFFs deviate slightly from the product rule, and this deviation is defined by ${}^{x,y}\gamma$ (24):

$$ {}^{x,y}\gamma = \frac{{}^{x,y}\alpha}{{}^{x}\alpha \times {}^{y}\alpha}, \tag{S4} $$

where $x$ and $y$ are the mass of the heavy isotopes. The $\gamma$ values are typically small and close to unity (25). We drew primary and secondary $\gamma$ values ($\gamma_p$ and $\gamma_s$, respectively) from uniform distributions:

$^{13,2}\gamma_p \in (0.998, 1)$ and $^{13,2}\gamma_s \in (1, 1)$ for $^{13}$C-D clumping, and $^{2,2}\gamma_p \in (0.994, 1)$ and $^{2,2}\gamma_s \in (1, 1)$ for double D clumping, which are similar to ranges that were explored in previous works (26).

### Data for model calibration and validation

#### Model calibration to laboratory culture results

We curated net carbon and hydrogen isotopic fractionations and their relations to the Gibbs free energy of the net methanogenic reaction ($\Delta G_{net}$) from culture experiments. Among other experimental parameters, this dataset consists of experiments with varying temperature, growth phases, substrates used and microbial strains. We screened the data according to the following criteria: (*i*) including data from stationary growth phases where growth was reported; (*ii*) excluding data that may have been affected by Rayleigh (isotopic) distillation; (*iii*) excluding data that were ignored in the original publications due to known measurement errors. For each sample we used the reported $\Delta G_{net}$, where available, or calculated it based on the reported concentrations of $H_2$, $CO_2$ and $CH_4$ using: $\Delta G_{net} = \Delta G_i'^0 + RT\ln Q$, where $\Delta G_i'^0$ is the standard Gibbs free energy of the reaction (corrected for the temperature with the Van't Hoff equation), R is the ideal gas constant, T is the temperature and $Q = [CH_4]/\left([CO_2][H_2]^4\right)$ is the reaction quotient. Isotopic data from experiments with small negative $\Delta G_{net}$ values were mostly obtained from co-cultures, which complicated measurements of headspace $H_2$ concentrations. We corrected for the concentrations of $H_2$ in these samples as explained above. All the samples that were used are listed in table S7.

The $^2\varepsilon_{CH_4\text{-}H_2O}$-$\Delta G_{net}$ relation at small negative $\Delta G_{net}$ values shows contradicting trajectories between two experiments that used a similar culture setup (3, 27). While both reports used the same thermophilic co-culture of *S. lipocalidus* and *M. thermoautotrophicus* strain ΔH, $^2\varepsilon_{CH_4\text{-}H_2O}$ values differ by $\sim$100‰ at similar $\Delta G_{net}$ values (fig. S15). As there were no physical differences between the two setups, for internal consistency we chose to calibrate our model against the results reported by Okumura et al. (3), which include measurements of both carbon and hydrogen isotopes over a larger range of $\Delta G_{net}$ values.

There are considerable differences between $\Delta^{13}CH_3D$ values in hydrogenotrophic methanogens with and without membrane-embedded electron carriers (e.g., methanophenazine), the latter of which are the focus of our model. While $^{2}\varepsilon_{CH_4\text{-}H_2O}$ is of a similar range for these two groups, $\Delta^{13}CH_3D$ for methanogens with methanophenazines is between –6 and 0‰, whereas for methanogens without methanophenazines the observed range is –2 to 3‰. The laboratory culture data collected to date is insufficient to determine whether this difference is statistically significant, though there may be a physiological basis for it, as methanogens with methanophenazines have distinct metabolic characteristics that may affect the dynamics of departure from equilibrium.

#### Comparison of isotopic fractionations between model and natural environments

We curated isotopic compositions of $CH_4$, $H_2O$ and $CO_2$ for both bulk isotopes and clumped isotopologues from various natural environments, focusing on samples of a known primary microbial origin and minimal indications of methane cycling (e.g., samples that are below the sulfate-methane transition zone in marine sediments). We divided the samples into categories: (*i*) marine sediments; (*ii*) coal-bed methane, shale gas and biogenic gas fields; (*iii*) brackish sediments; and (*iv*) enrichment cultures (fig. S16). A list of the data and the different categories that were assigned to them and their references is in table S8.

#### Cell-specific methanogenesis rates

There are currently limited data on cell-specific methanogenesis rates in natural environments. We bridge this gap by comparing compiled bulk methanogenesis rates (bMR) and estimates of cell density. bMR values were obtained from the results of either ratiotracer experiments or reaction-diffusion models, both of which carry uncertainties. In radiotracer experiments, the rate of methanogenesis is assumed to be equal to the rate of $CO_2$ reduction to methane, but in fact the measured rate of $CO_2$ reduction serves as an upper limit on methanogenesis rates. If the reaction is close to equilibrium, then the net rate of methanogenesis will be lower than the radiotracer-based estimate, possibly by orders of magnitude if the reversibility between methane and $CO_2$ is higher than 0.9. Moreover, some of the radiotracer experiments are conducted under conditions that may favor higher methanogenesis rates (e.g., increased partial pressure of $H_2$ in the headspace), resulting in overestimation of the in-situ rates. Models of in-situ methanogenesis rates provide an estimate for the net methanogenesis rates based on the concentration and isotopic gradients of methane and DIC, but carry uncertainties due to the choice of model parameters, such as the net fractionation of carbon and hydrogen isotopes associated with methanogenesis.

To estimate the abundance of cells, where no measurements exist, we used a general relation between cell density and depth within the sediment in marine environments (28, 29): $y = 7.73 \cdot 10^7 \times z^{-0.6332}$, where $z$ is the depth in meters and $y$ is the number of cells per cubic centimeter of sediment. We assume that of these cells 12% are Archaea in open-ocean sites and 40% in ocean margin sites (30), and that 50% of Archaea are methanogens (29).

### Supplementary Figures

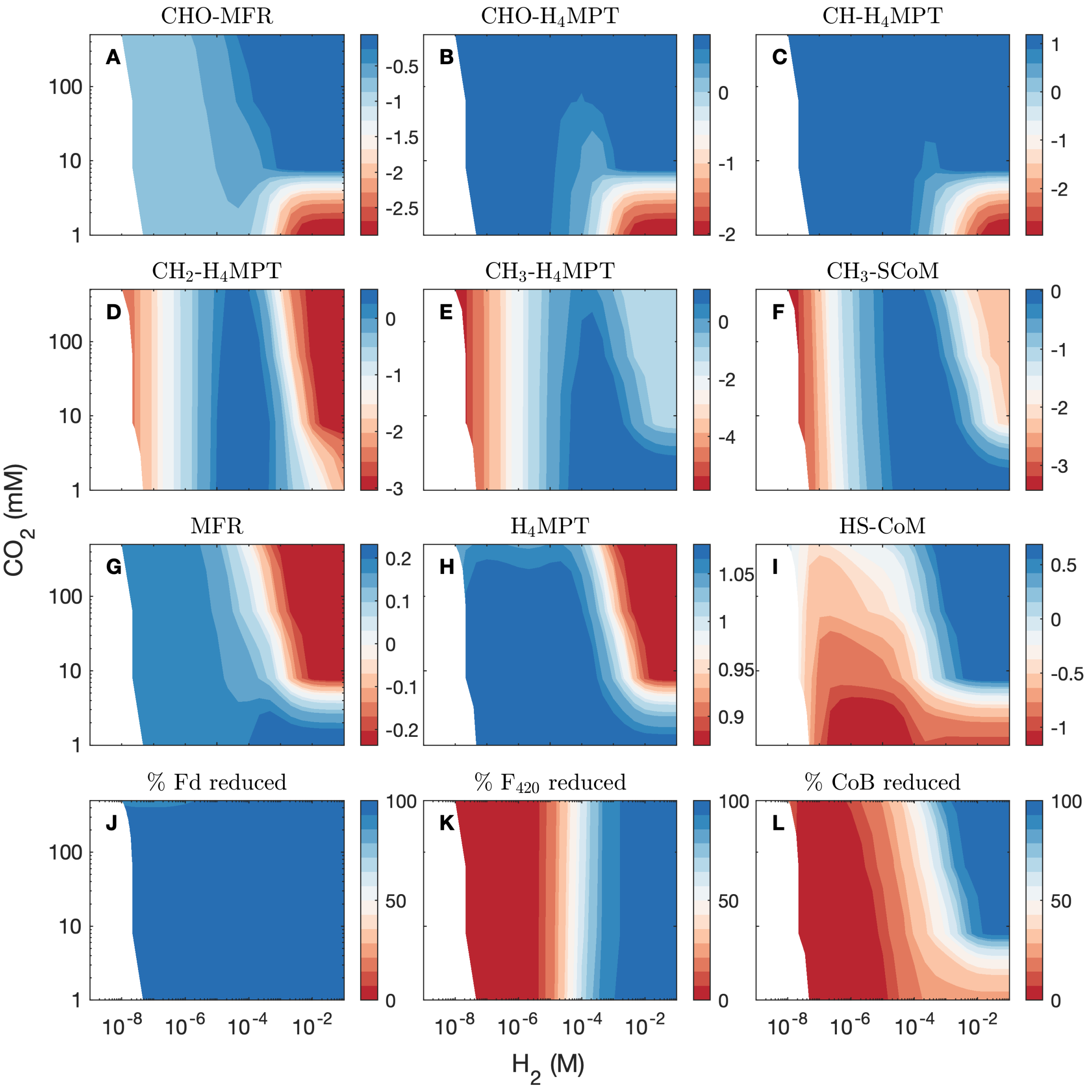


**Figure S1: Intracellular metabolite concentrations.** All simulations are at [$CH_4$] of 0.01 mM and 60°C, under Mcr II conditions. (**A**-**I**) $\log_{10}$ of concentrations in M (see individual panel color legends). (**J**-**L**) Percentage of electron carriers in reduced form.

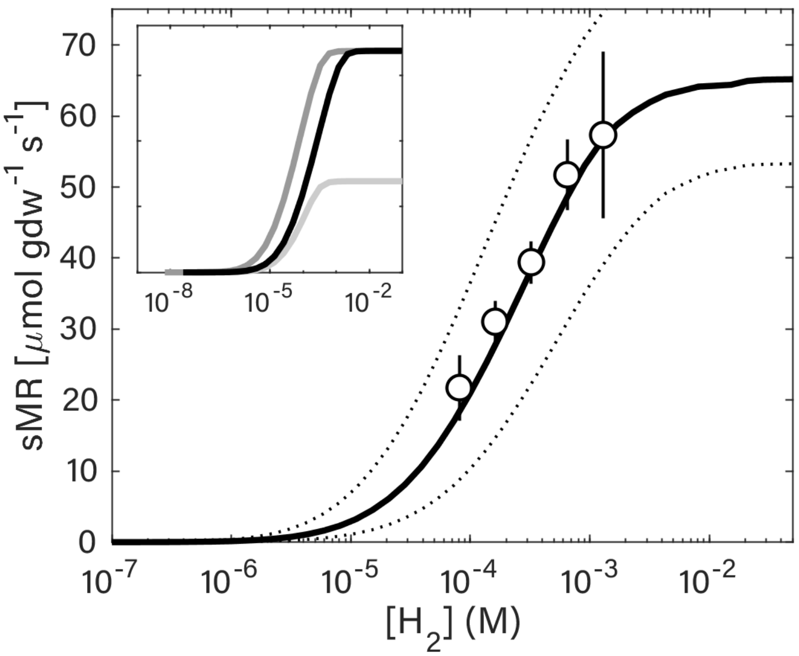


**Figure S2: Specific methanogenesis rates (sMR) depend on the $H_2$ concentration.** Measured sMR (circles) during the linear and stationary phase in laboratory cultures of *M. thermoautotrophicus* grown at 60°C (error bars represent 1$\sigma$ of the results of four experiments) (11). The results of $10^6$ model simulations are shown (median and 95% of model results in the solid and dotted lines, respectively). Metabolic model parameter combinations were drawn from posterior distributions resulting from the model calibration (see SM text and fig. S14). The inset shows the same results for [$CO_2$] of 0.1 M (dark grey) and 0.005 M (light grey) in addition to the default 0.01 M (black).

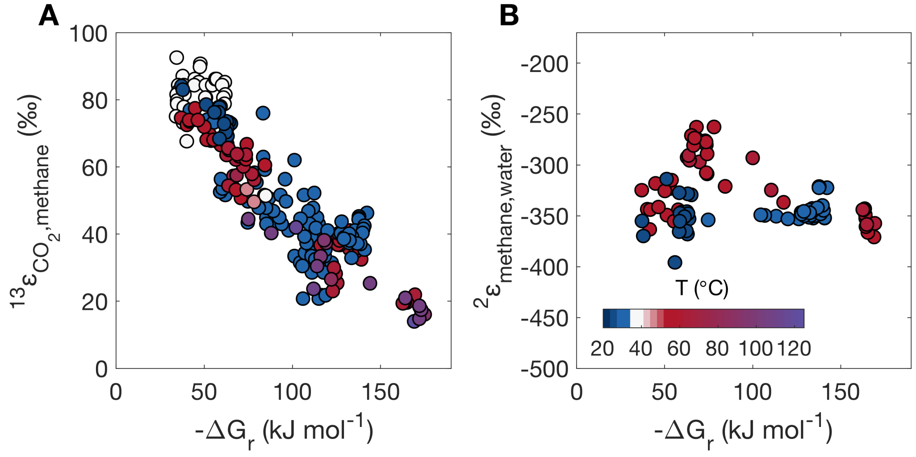


**Figure S3: Isotopic fractionation in laboratory cultures depends on the thermodynamic drive.** A compilation of (**A**) carbon isotopic fractionation between $CO_2$ and $CH_4$ ($^{13}\varepsilon_{CO_2\text{-}CH_4}$), and (**B**) the hydrogen isotopic fractionation between $CH_4$ and $H_2O$ ($^{2}\varepsilon_{CH_4\text{-}H_2O}$). The colors correspond to the temperature of the culture as shown in the color bar in panel **B**. The data were taken from (2, 3, 27, 31–34), and are listed in table S7.

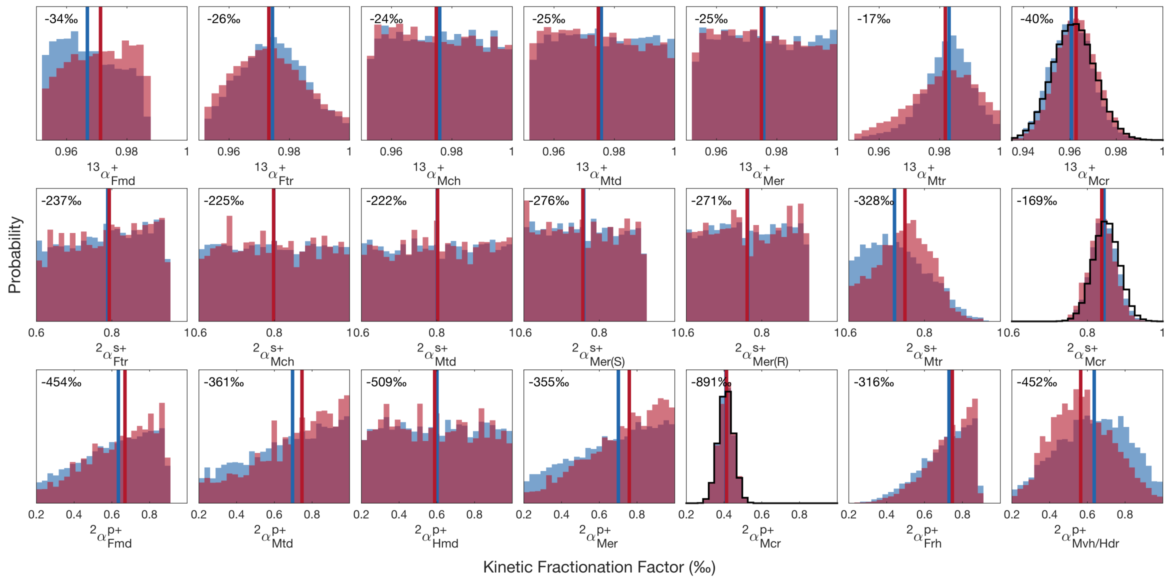


**Figure S4: Posterior distributions of kinetic isotope fractionation factors (KFFs).** The posterior KFF distributions were generated by weighting individual KFF values by the inverse of the square of sum of squared errors ($1/SSE^2$) of $1 \times 10^6$ simulations from the prior distributions, for mesophilic (blue) and thermophilic (red) conditions. Carbon KFFs (top row) were drawn from uniform prior distributions with $\alpha \in (0.95, 1)$. The KFF of Mcr was assigned a normal prior distribution with $0.952 \pm 0.01$, as constrained by experiments (20). Secondary hydrogen KFFs (middle row) were drawn from uniform prior distributions with $\alpha \in (0.6, 1)$, and primary hydrogen KFFs (bottom row) were drawn from prior uniform distributions with $\alpha \in (0.2, 1)$. The primary and secondary hydrogen KFFs of Mcr were drawn from normal prior distributions of $0.41 \pm 0.04$ and $0.85 \pm 0.0035$, respectively (20). The vertical lines in each panel represent the median KFF value that was used to generate the median model results. The black histogram outlines in the KFFs of Mcr represent the prior (non-uniform) distributions. Secondary and primary hydrogen KFFs are denoted by superscripted 'p' and 's' respectively.

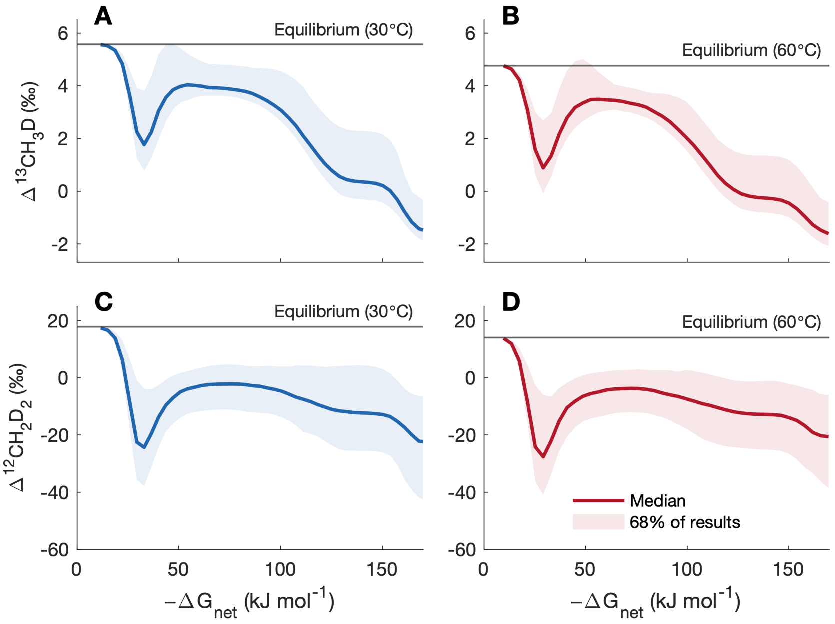


**Figure S5: Clumped isotopologue abundances.** Model results of 200 simulations at 30°C (blue) and 60°C (red) against $\Delta G_{net}$. The gray lines represent temperature-dependent isotopic equilibrium. (**A-B**) $\Delta^{13}CH_3D$. (**C-D**) $\Delta^{12}CH_2D_2$.

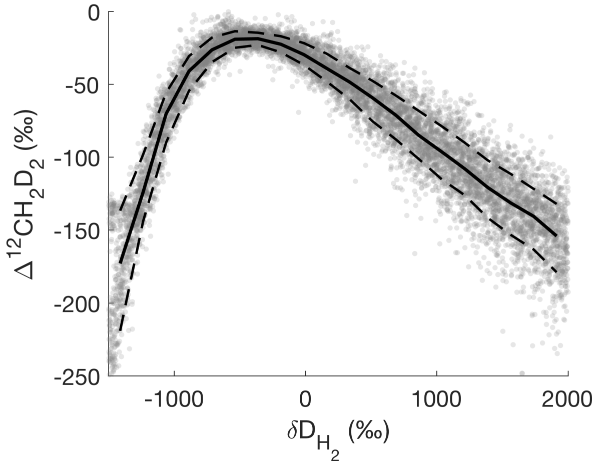


**Figure S6: A combinatorial effect in hydrogenotrophic methanogenesis due to activity of Hmd.** Methane $\Delta^{12}CH_2D_2$ values against $H_2$ $\delta D$ values calculated at 60°C, with $H_2$ and $CO_2$ concentrations of 50 mM and a $CH_4$ concentration of 10 $\mu$M. The $\delta D$ values of $H_2O$ in all the calculations are 0‰. The solid line is the median of $10^4$ simulations, and the dashed envelope contains 68% of the results. The variability in the $\Delta^{12}CH_2D_2$ values is due to the hydrogen KFFs, which were randomly drawn from the posterior distributions presented in fig. S4, with constant KFFs for Hmd (0.95) and Mvh/Hdr (0.45).

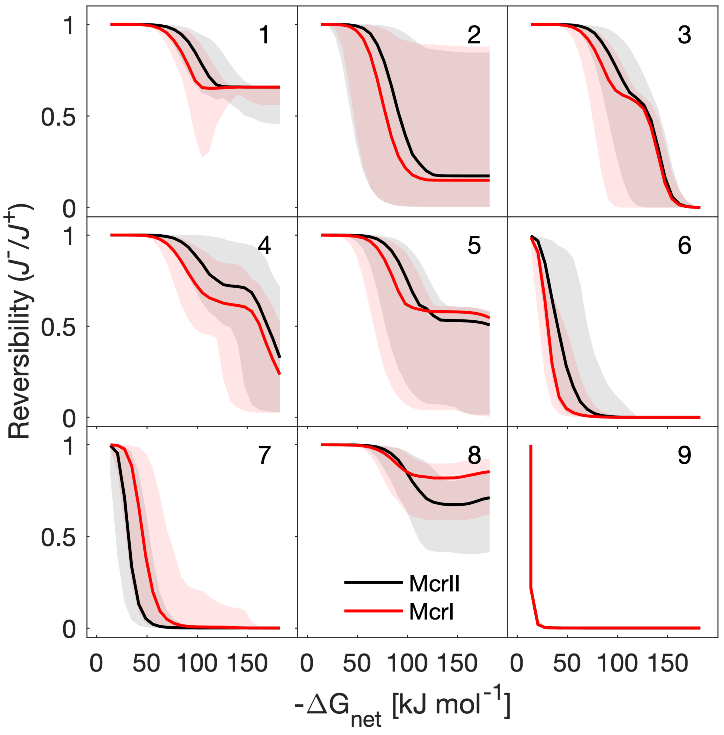


**Figure S7: Reversibility against the thermodynamic drive ($\Delta G_{net}$) of methanogenesis under energy-limiting conditions.** Mcr II results simulate growth under optimal conditions in lab cultures (red), and Mcr I is for growth in energy-limited conditions (black). The $\Delta G_{net}$ is for $H_{2(aq)}$ concentrations in the range 1 nM to 10 mM, $CO_2$ = 10 mM and $CH_4$ = 10 $\mu$M, which represent typical experimental conditions. The subplot titles correspond to the reactions in table S1. The solid lines are the median of 100 model simulations, and the envelopes represent 95% of model results. The uncertainty originates mostly from the kinetic parameters of the enzymatically-catalyzed reactions. The reversibility, defined as the ratio of the backward to forward reaction rates of an individual reaction ($J_i^-/J_i^+$) ranges from 1 (full reversibility) to 0 (unidirectional reaction).

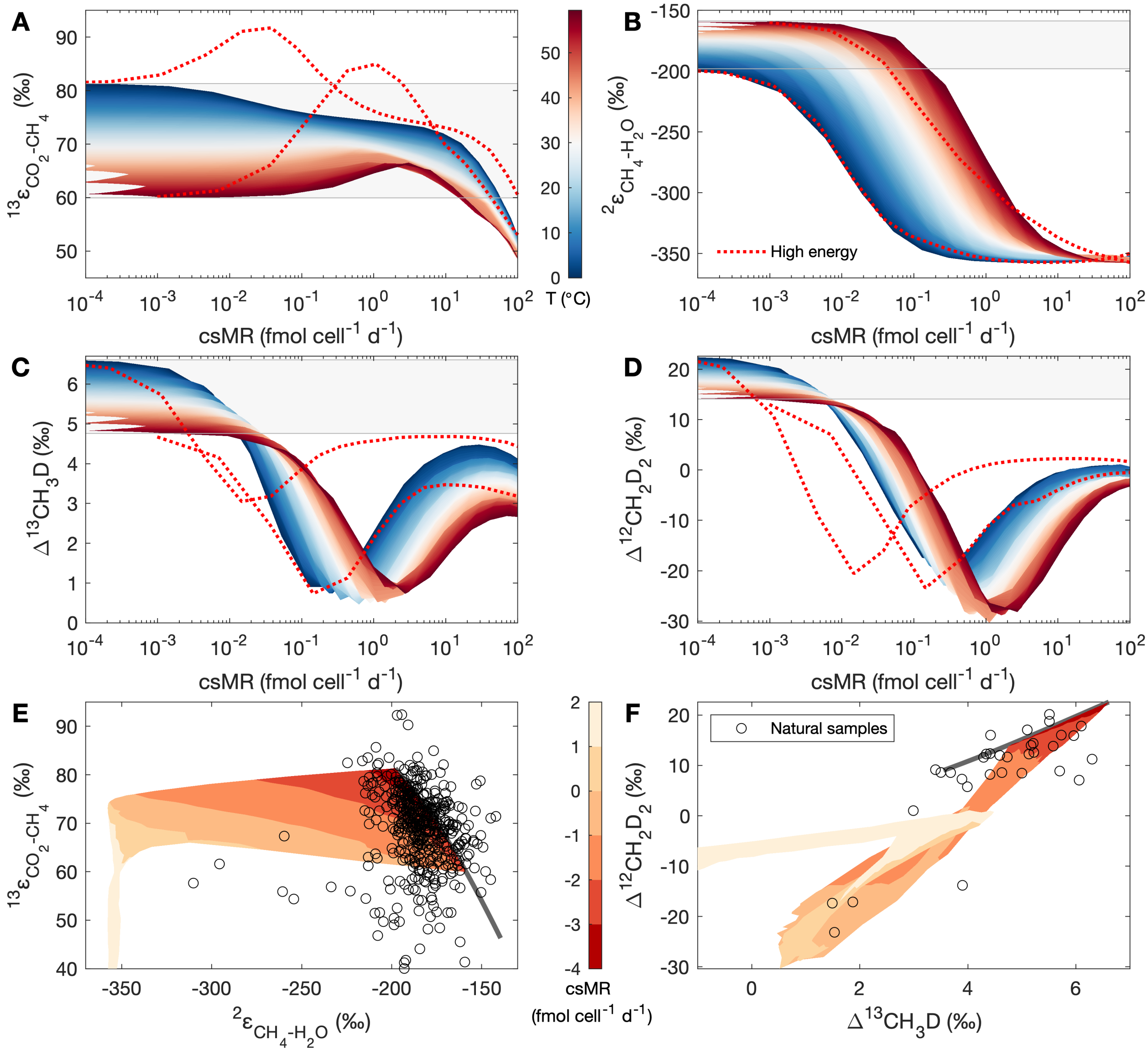


**Figure S8: Isotopic fractionation during methanogenesis in energy-limited conditions.** The dependence on csMR of (**A**) $^{13}\varepsilon_{CO_2\text{-}CH_4}$, (**B**) $^{2}\varepsilon_{CH_4\text{-}H_2O}$, (**C**) $\Delta^{13}CH_3D$, and (**D**) $\Delta^{12}CH_2D_2$. The light-gray envelopes in panels **A-D** represent the equilibrium isotopic fractionations between 0°C and 60°C. (**E**) Co-variation of $^{13}\varepsilon_{CO_2\text{-}CH_4}$ and $^{2}\varepsilon_{CH_4\text{-}H_2O}$. (**F**) Co-variation of $\Delta^{12}CH_2D_2$ and $\Delta^{13}CH_3D$. The contours in panels **E-F** are the $\log_{10}$ of the csMR as predicted by our metabolic model between 0°C and 60°C, and the black circles are biogenic environmental samples (table S8). In panels **A-D**, the dotted red lines show the default laboratory-calibrated model results for the same temperatures (Mcr II). The calculations are for a $H_2$ concentration range of 1 nM to 5 $\mu$M, and $CO_2$ and $CH_4$ concentrations of 1 mM. The cell volume is 1 $\mu m^3$.

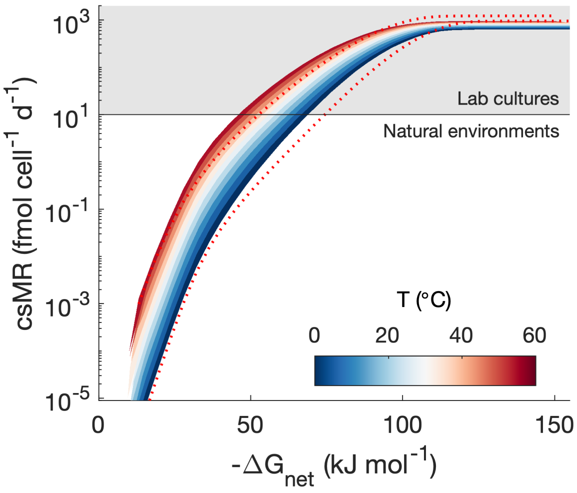


**Figure S9: The dependence of cell-specific methanogenesis rates (csMRs) on $\Delta G_{net}$.** The dotted red lines show the default laboratory-calibrated model results for the same temperatures. The calculations are for a $H_2$ concentration range of 1 nM to 5 $\mu$M, and $CO_2$ and $CH_4$ concentrations of 1 mM. The cell volume is 1 $\mu m^3$.

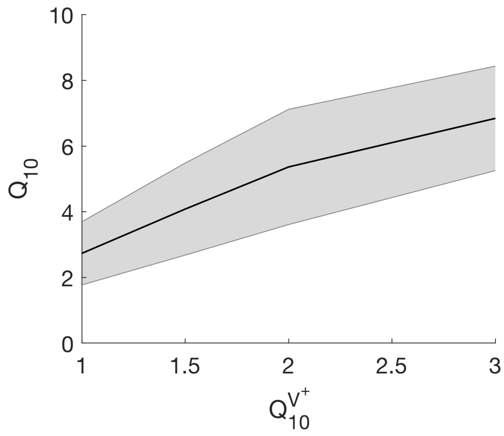


**Figure S10: The temperature coefficient ($Q_{10}$) of methanogenesis.** The dependence of $Q_{10}$ on the scaling factor of metabolic rate capacity ($Q_{10}^{V^+}$). The gray envelope represents $1\sigma$ of the results for a $\Delta G_{net}$ range between –10 to –60 $\mathrm{kJ\,mol^{-1}}$. All the simulations are for $H_2$ concentrations of 1 to 1000 nm, and $CO_2$ and $CH_4$ concentrations of 1 mM, and under energy-limited conditions (Mcr I).

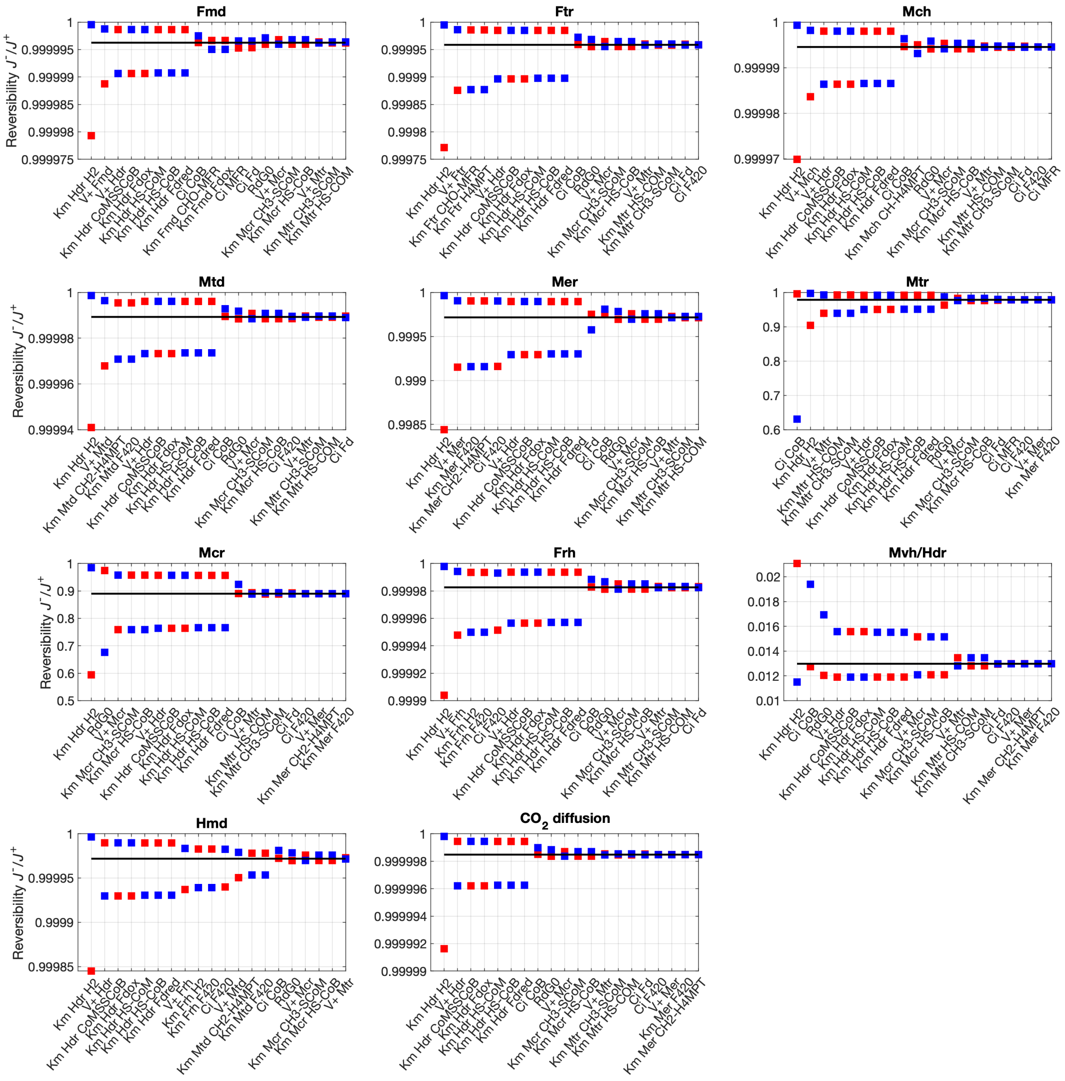


**Figure S11: Sensitivity of modeled reversibility to enzyme kinetic parameters at $H_2$ of 50 nM and 60°C.** Equivalent to $\Delta G_r$ of −22 kJ mol$^{-1}$, with $CO_2$ of 10 mM and $CH_4$ of 10$\mu$M. In each panel, we present the 20 parameters that had the largest effect over the reversibility $J^-/J^+$. Black lines represent results obtained with the default parameters, red and blue squares represent an increase or a decrease by 1/3 of the default parameter values, respectively. Note that the y-axis scales are not aligned to the full range of possible $J^-/J^+$ values, and in some cases cover a very small range around the default result.

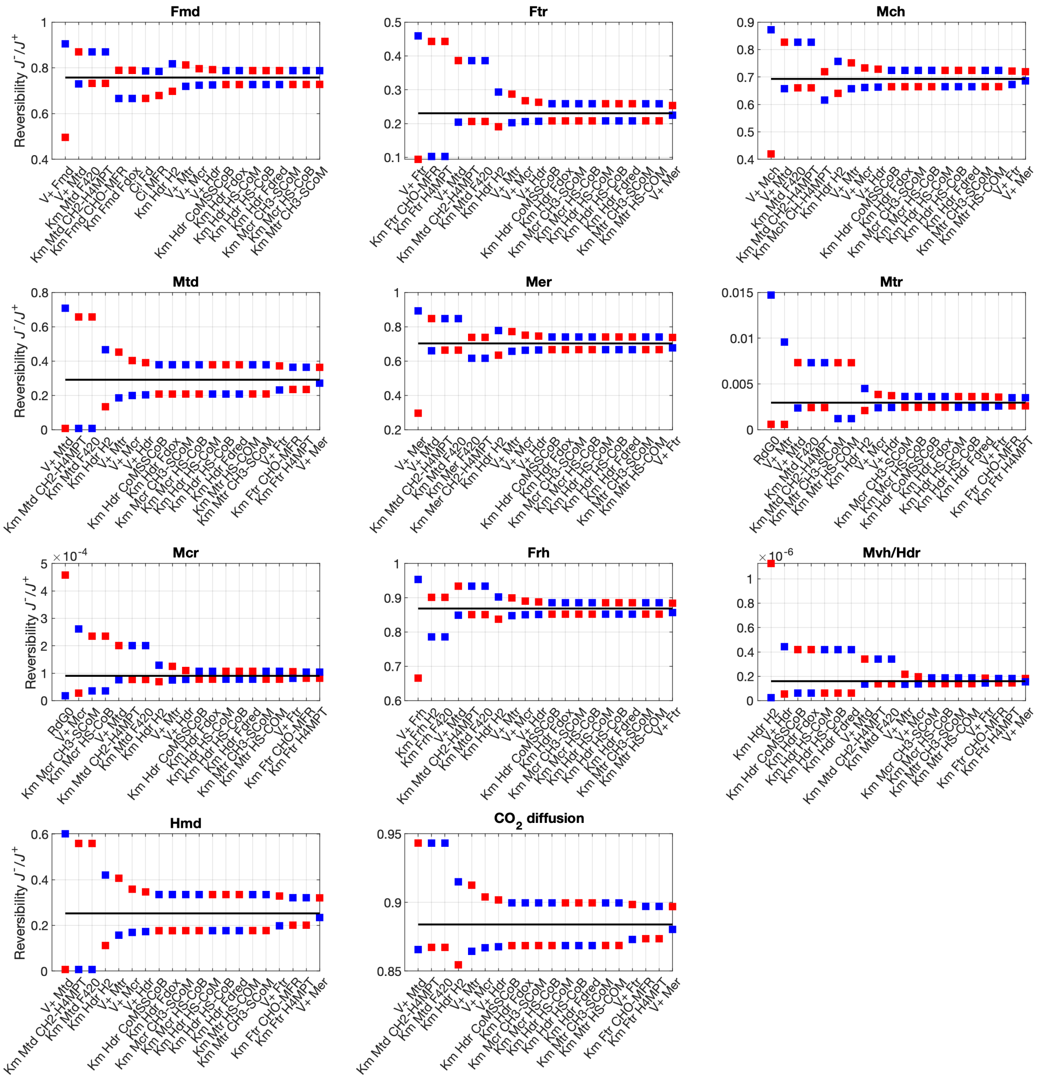


**Figure S12: Sensitivity of modeled reversibility to enzyme kinetic parameters at $H_2$ of 100 $\mu$M and 60**°C**.** Equivalent to $\Delta G_r$ of –106 kJ mol$^{-1}$, with $CO_2$ of 10 mM and $CH_4$ of 10$\mu$M. In each panel, we present the 20 parameters that had the largest effect over the reversibility $J^-/J^+$. Black lines represent results obtained with the default parameters, red and blue squares represent an increase or a decrease by 1/3 of the default parameter, respectively. Note that the y-axis scales are not aligned to the full range of possible $J^-/J^+$ values, and in some cases cover a very small range around the default result.

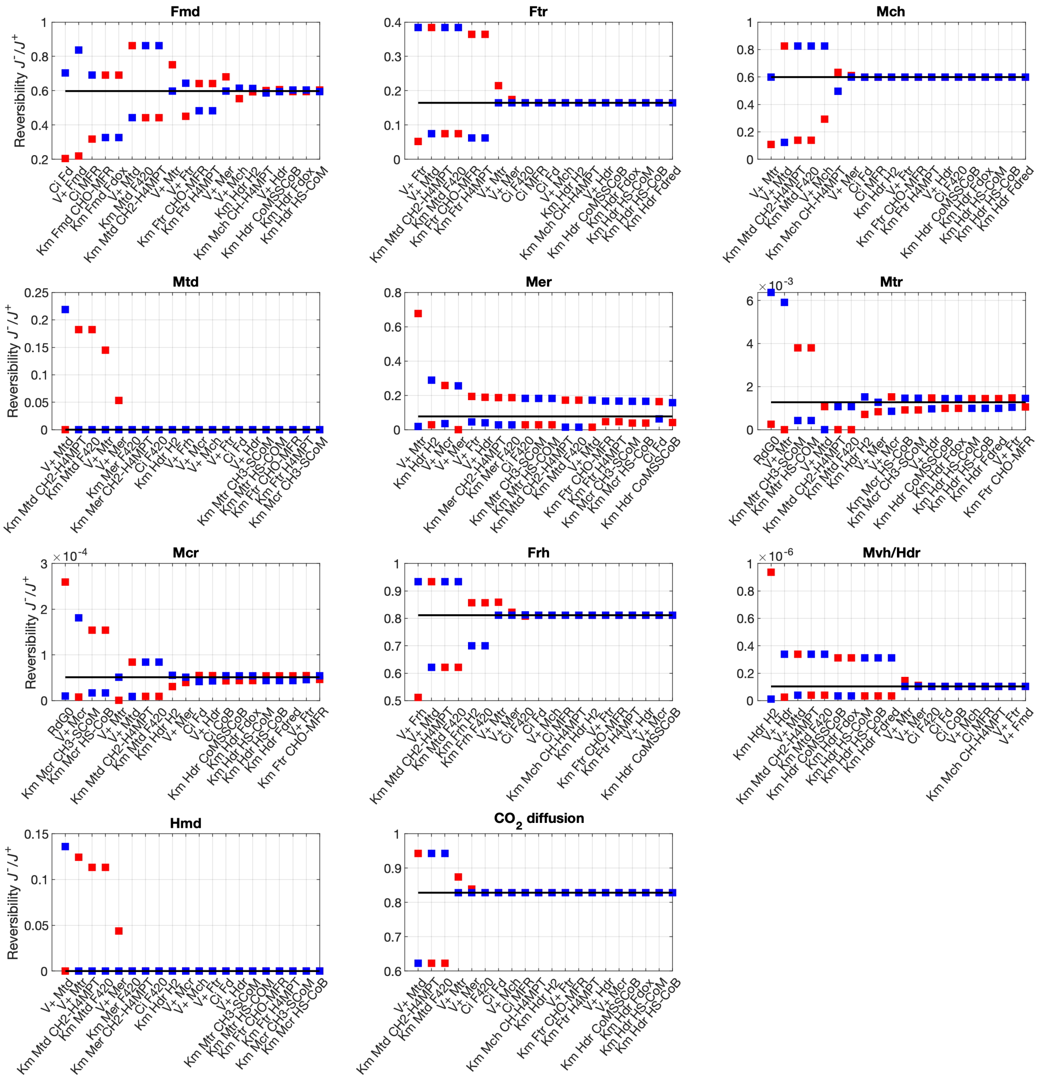


**Figure S13: Sensitivity of modeled reversibility to enzyme kinetic parameters at $H_2$ of 10 mM and 60°C.** Equivalent to $\Delta G_r$ of –157 kJ mol$^{-1}$, with $CO_2$ of 10 mM and $CH_4$ of 10$\mu$M. In each panel, we present the 20 parameters that had the largest effect over the reversibility $J^-/J^+$. Black lines represent results obtained with the default parameters, red and blue squares represent an increase or a decrease by 1/3 of the default parameter, respectively. Note that the y-axis scale are not aligned to the full range of possible $J^-/J^+$ values, and in some cases cover a very small range around the default result.

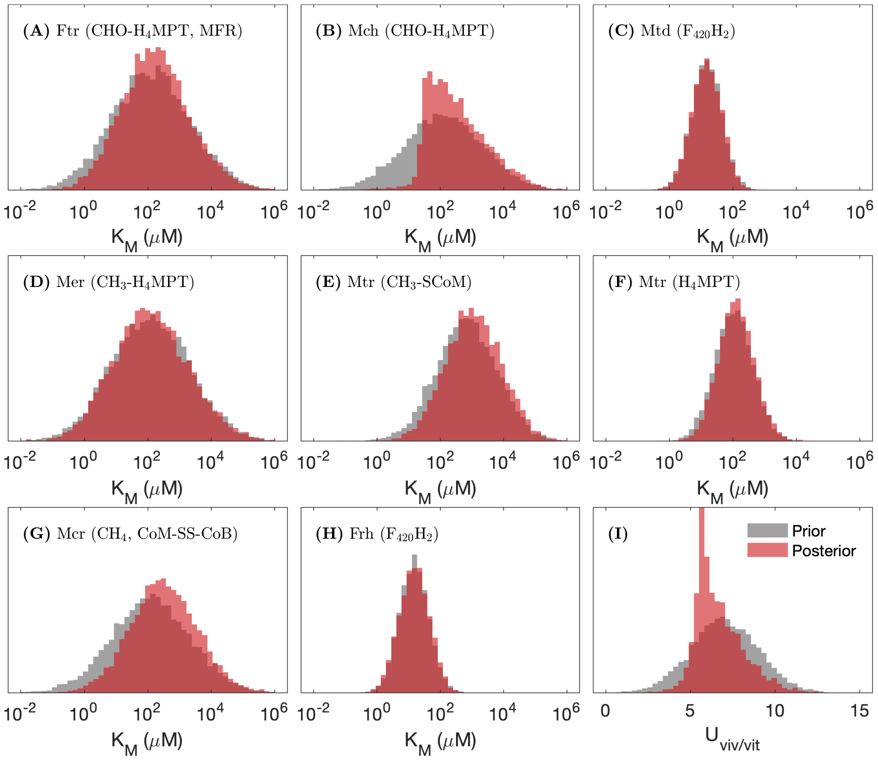


**Figure S14: Prior and posterior distributions of metabolic model parameters.** (**A–H**) $K_m$ values and (**I**) $U_{(viv/vit)}$, for $10^6$ simulations. Posterior distributions that differ markedly from the prior distributions indicate sensitivity to the model parameter.

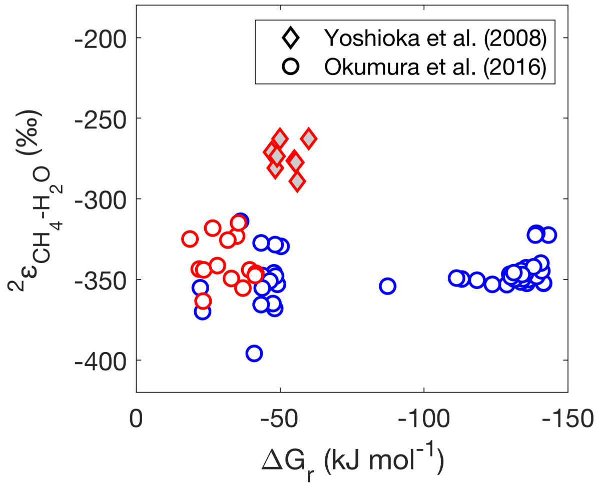


**Figure S15: Hydrogen isotopic fractionation in laboratory cultures.** Laboratory cultures of methanogens show a variable relation between $^2\varepsilon_{CH_4\text{-}H_2O}$ and $\Delta G_r$, for mesophilic (blue symbols) and thermophilic (red symbols) conditions. Both reports used an identical syntrophic co-culture of a butyrate oxidizing bacteria (*Syntrophothermus lipocalidus*, DSMZ 12681) and a thermophilic methanogen (*Methanothermobacter thermautotrophicus* strain ΔH JCM10044). However, they yield different $^2\varepsilon_{CH_4\text{-}H_2O}$ values at small negative $\Delta G_r$ probably due to different measurement techniques (3, 27). We chose to calibrate our model relative to the results from the experiment by (3) (circles), for internal consistency with the measurements of carbon and hydrogen isotopes in mesophilic conditions.

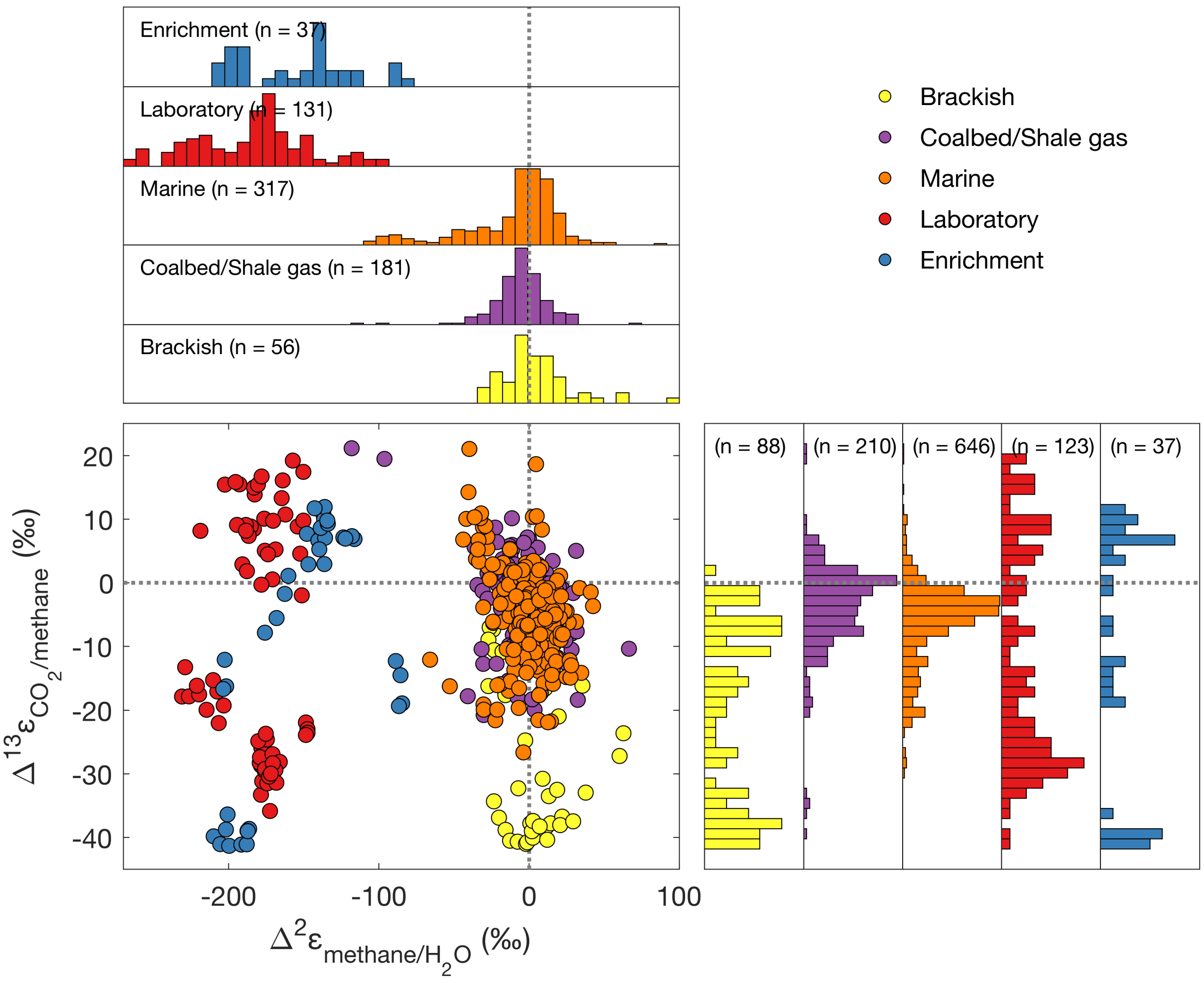


**Figure S16: Deviation of apparent isotopic fractionation from equilibrium.** A compilation of apparent carbon and hydrogen isotopic fractionations from the temperature-dependent equilibrium isotopic fractionation, from laboratory culture experiments and natural environments. A list of the samples that are included here is in table S8. We included samples of known biogenic origin from natural environments, below sulfate-methane interfaces, where these exist. We did not include here samples from terrestrial sources, which are typically a mixture of acetoclastic and hydrogenotrophic methanogenic activity. The dotted lines represent the equilibrium isotopic fractionation at a given temperature (i.e., samples at the crossover of these lines are in both carbon and hydrogen isotopic equilibrium). The numbers of samples from each group are shown in parentheses (not all samples have information on both carbon and hydrogen isotopes).

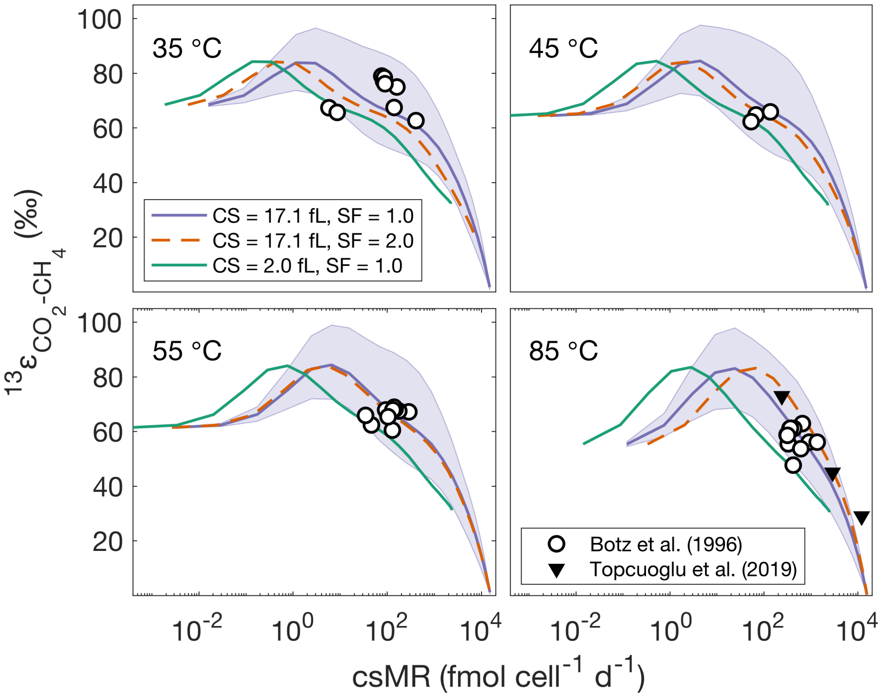


**Figure S17: The dependence of carbon isotope fractionation on cell-specific methanogenesis rates (csMR).** The carbon isotope fractionation between $CO_2$ and $CH_4$ ($^{13}\varepsilon_{CO_2\text{-}CH_4}$) was calculated at 35°C, 45°C, 55°C and 85°C. In panel **A**, the legend is for different cell-sizes (CS) and $Q_{10}^{V+}$ values (SF). Am optimal fit was obtained with a cell volume of 17.1 fL, and a $Q_{10}^{V+}$ of 1. The solid lines are the median value of the model and the envelopes include 95% of the model results from 1000 simulations using the posterior distributions of the carbon isotope KFFs. Concentrations of $CO_2$ and $CH_4$ are 10 mM and 10 $\mu$M, respectively. We used experimental data from cultures in the stationary growth phase (14, 34).

### Supplementary Tables

**Table S1:** Transformed Gibbs free energy of the reactions in hydrogenotrophic methanogenesis. Whenever possible, we used the transformed Gibbs free energy ($\Delta G_i'^0$) at pH = 7 (35). For redox reactions, we calculated $\Delta G_i'^0$ of the half-reactions using the Nernst equation with $\Delta G_i'^0 = -nFE'$, where n is the numbers of electrons transferred, F is the Faraday constant and $E'$ is the standard redox potential.

| | Enzyme | Reaction | | | $\Delta G_r'^0$ ($kJ\,mol^{-1}$) | Ref./Source |
|---|---|---|---|---|---|---|
| | | $4H_{2(aq)} + CO_{2(aq)}$ | $\rightleftharpoons$ | $CH_{4(aq)} + 2H_2O$ | –188.8 | † |
| | | $CO_{2(out)}$ | $\rightleftharpoons$ | $CO_{2(in)}$ | | |
| 1 | Fmd | $CO_{2(aq)}$ + MFR + $Fd^{2-}_{red}$ | $\rightleftharpoons$ | CHO-MFR + $Fd_{ox}$ | 10.3 | (36, 37) |
| 2 | Ftr | CHO-MFR + $H_4$MPT | $\rightleftharpoons$ | CHO-$H_4$MPT + MFR | –3.5 | (38) |
| 3 | Mch | CHO-$H_4$MPT | $\rightleftharpoons$ | CH≡$H_4MPT^+$ + $H_2O$ | –4.2 | (38) |
| 4a | Mtd | CH≡$H_4MPT^+$ + $F_{420}H_2$ | $\rightleftharpoons$ | $CH_2$=$H_4$MPT + $F_{420}$ | 2.0 | (38) |
| 4b | Hmd | CH≡$H_4MPT^+$ + $H_{2(aq)}$ | $\rightleftharpoons$ | $CH_2$=$H_4$MPT | –27.6 | § |
| 5 | Mer | $CH_2$=$H_4$MPT + $F_{420}H_2$ | $\rightleftharpoons$ | $CH_3$-$H_4$MPT + $F_{420}$ | –1.7 | (38) |
| 6 | Mtr | $CH_3$-$H_4$MPT + HS-CoM | $\rightleftharpoons$ | $CH_3$-S-CoM + $H_4$MPT | –31.4 | ¶ |
| 7 | Mcr | $CH_3$-S-CoM + HS-CoB | $\rightleftharpoons$ | $CH_{4(aq)}$ + CoM-S-S-CoB | –13.5 | ¶ |
| 8 | Frh | $F_{420}$ + $H_{2(aq)}$ | $\rightleftharpoons$ | $F_{420}H_2$ | –28.0 | (39) |
| 9 | Mvh/Hdr | $2H_{2(aq)}$ + CoM-S-S-CoB + $Fd_{ox}$ | $\rightleftharpoons$ | HS-CoM + HS-CoB + $Fd^{2-}_{red}$ | –90.8 | (36, 40) |
| | ATP | AFP + Pi + $4Na^+_{(out)}$ | $\rightleftharpoons$ | ATP + $4Na^+_{(in)}$ | 46.2 | (9) |

Abbreviations: $F_{420}$, coenzyme $F_{420}$; Fd, ferredoxin; $H_4$MPT, tetrahydromethanopterin; HS-CoB, coenzyme B; HS-CoM, coenzyme M; MFR, methanofuran; Fmd, formyl-MFR dehydrogenase; Ftr, formyltransferase; Mch, methenyl cyclohydrolase; Mtd, methylene-$H_4$MPT dehydrogenase; Hmd, $H_2$-dependent methylene-$H_4$MPT dehydrogenase; Mer, methylene-$H_4$MPT reductase; Mtr, methyl transferase, Mcr, methyl-CoM reductase; Frh, $F_{420}$ reducing hydrogenase; Mvh/Hdr, methylviologen hydrogenase/heterodisulfide reductase.

† Net methanogenesis at 60°C, calculated by Van't Hoff's Equation. Whenever possible, $\Delta G_i'^0$ for individual reactions were at 60°C.

§ Internally consistent value, inferred from summation of reactions 4 and 8 in this table.

¶ Reactions 6 and 7 have an estimated $\Delta G_i'^0$ of –30±10 $kJ\,mol^{-1}$ each (with $CH_4$ in the gaseous phase). We allocate the exact $\Delta G_i'^0$ of reactions 6 and 7 as the remaining $\Delta G_i'^0$ from the $\Delta G_i'^0$ of reactions 1–5 and 8–9 and the total pathway.

**Table S2:** Metabolic model kinetic parameters.

| $V^+$ ($\mu$mol gdw$^{-1}$ s$^{-1}$) | Metabolite | $K_M$ (mM) | Ref/Source | Notes |
|---|---|---|---|---|
| **1. formyl-MFR dehydrogenase (Fmd): $CO_2$ + MFR + $Fd_{red}$ ⇄ CHO-MFR + $Fd_{ox}$** | | | | |
| 10.96 | | | (41) | Inferred from $V^+/V^- = 25$ |
| | $CO_2$, MFR, $Fd_{red}$ | 6.800 | (36) | |
| | CHO-MFR | 0.030 | (41, 42) | Harmonic mean of two values |
| | $Fd_{ox}$ | 0.010 | | Estimate based on methylviologen as electron carrier |
| **2. formyl transferase (Ftr): CHO-MFR + $H_4$MPT ⇄ CHO-$H_4$MPT + MFR** | | | | |
| 51.35 | | | (43) | |
| | CHO-MFR | 0.050 | (44) | Harmonic mean of two values |
| | $H_4$MPT | 0.060 | (44) | Estimate based on *M. barkeri* |
| | CHO-$H_4$MPT, MFR | 0.071 | ‡ | |
| **3. methenyl cyclohydrolase (Mch): CHO-$H_4$MPT ⇄ CH-$H_4$MPT** | | | | |
| 7.26 | | | (45) | |
| | CHO-$H_4$MPT | 0.250 | ‡ | |
| | CH-$H_4$MPT | 0.167 | (46, 47) | Estimate based harmonic mean of two values from *M. barkeri* and *M. kandleri* |
| **4a. methylene dehydrogenase (Mtd): CH-$H_4$MPT + $F_{420}H_2$ ⇄ $CH_2$-$H_4$MPT + $F_{420}$** | | | | |
| 26.45 | | | (48, 49) | Harmonic mean of two values |
| | CH-$H_4$MPT | 0.050 | (50) | |
| | $F_{420}H_2$ | 0.028 | ‡ | |
| | $CH_2$-$H_4$MPT | 0.033 | (49) | |
| | $F_{420}$ | 0.065 | (49) | |
| **4b. $H_2$-producing methylene dehydrogenase (Hmd): CH-$H_4$MPT + $H_2$ ⇄ $CH_2$-$H_4$MPT** | | | | |
| 82.17 | | | (51) | |
| | CH-$H_4$MPT | 0.050 | (50) | |
| | $H_2$ | 0.150 | (52) | |
| | $CH_2$-$H_4$MPT | 0.040 | (50) | |
| **5: methylene reductase (Mer): $CH_2$-$H_4$MPT + $F_{420}H_2$ ⇄ $CH_3$-$H_4$MPT + $F_4$20** | | | | |
| 10.96 | | | (53) | |
| | $CH_2$-$H_4$MPT | 0.300 | (53) | Estimate based on *M. marburgensis* |
| | $F_{420}H_2$ | 0.003 | (53) | Estimate based on *M. marburgensis* |
| | $CH_3$-$H_4$MPT | 0.092 | ‡ | |
| | $F_{420}$ | 0.040 | (54) | Estimate based on *M. barkeri* |

Table continued on next page.

| $V^+$ ($\mu$mol gdw$^{-1}$ s$^{-1}$) | **Metabolite** | $K_M$ (mM) | **Ref/Source** | **Notes** |
|---|---|---|---|---|
| **6. methyl transferase (Mtr): $CH_3$-$H_4$MPT + HS-CoM ⇄ $CH_3$-S-CoM + $H_4$MPT** | | | | |
| 8.55 | | | (6) | Original value measured at 37°C and was multiplied ×2.5 |
| | $CH_3$-$H_4$MPT | 0.135 | (55) | Estimate based on *M. acetivorans* |
| | HS-CoM | 0.277 | (55) | Estimate based on *M. acetivorans* |
| | $CH_3$-S-CoM | 0.504 | ‡ | |
| | $H_4$MPT | 0.094 | ‡ | |
| **7. methyl-CoM reductase (Mcr): $CH_3$-S-CoM + HS-CoB ⇄ $CH_4$ + CoM-S-S-CoB** | | | | |
| 18.73 (136.95†) | | | (11, 56, 57) | Harmonic mean of three values |
| | $CH_3$-S-CoM | 0.821 (0.280†) | (57, 58) | Harmonic mean of three values |
| | HS-CoB | 0.204 (0.075†) | (57, 59, 60) | Harmonic mean of three values |
| | $CH_4$, CoM-S-S-CoB | 0.082 | | |
| **8. $F_{420}$ hydrogenase (Frh): $F_{420}$ + $H_2$ ⇄ $F_{420}H_2$** | | | | |
| 65.75 (131.50†) | | | (11, 61) | Original value measured at 32°C and was multiplied by ×3 |
| | $H_2$ | 0.012 | (61) | |
| | $F_{420}$ | 0.036 | (61) | |
| | $F_{420}H_2$ | 0.014 | ‡ | |
| **9. methyl viologen hydrogenase/heterodisulfide reductase (Mvh/Hdr): CoM-S-S-CoB + $Fd_{ox}$ + 2$H_2$ ⇄ HS-CoM + HS-CoB + $Fd_{red}$** | | | | |
| 34.24 (102.72†) | | | (11, 62) | |
| | $H_2$ | 0.030 | (62, 63) | |
| | CoM-S-S-Co B | 0.145 | (63) | |
| | $Fd_{ox}$ | 0.010 | (64) | Estimate based on *M. barkeri* |
| | HS-CoM | 0.200 | (65) | |
| | HS-CoB | 0.200 | (65) | |
| | $Fd_{red}$ | 0.075 | (64) | |

‡ Median values drawn from the posterior distributions (see Section ).

† Values that were for simulating methanogenesis in $H_2$-limited conditions.

**Table S3:** Physical parameters used in the bio-isotopic model.

| Parameter | Value | Units | Ref. | Notes |
|---|---|---|---|---|
| Cell volume | 2 | fL/cell | (66) | *M. thermoautotrophicus* |
| Cell mass | $10^{-12}$ | gdw/cell | (67) | *M. thermoautotrophicus* |
| Dry weight protein content | 41.1 | % | (68) | *M. bryantii* |
| Membrane thickness | 0.5 | nm | – | |
| Diffusivity constant | $2.9\times10^{-9}$ | $m^2\ s^{-1}$ | (1) | |
| Cell radius | 0.5 | $\mu$m | – | |
| $K_H$ ($H_2$) | $6.46\times10^{-9}$ | mol $L^{-1}$ $Pa^{-1}$ | (69) | Henry's Law constant at 60°C |
| $K_H$ ($CO_2$) | $1.42\times10^{-7}$ | mol $L^{-1}$ $Pa^{-1}$ | (69) | Henry's Law constant at 60°C |
| $K_H$ ($CH_4$) | $7.97\times10^{-9}$ | mol $L^{-1}$ $Pa^{-1}$ | (69) | Henry's Law constant at 60°C |
| Total ferredoxin (Fd)† | 5 | mM | – | |
| Total coenzyme $F_{420}^{\dagger}$ | 0.5 | mM | (15, 68, 70) | |
| Total coenzyme B (HS-CoB)† | 6 | mM | (71) | |
| Total methanofuran (MFR)† | 1.8 | mM | (71) | |

† Total Fd and $F_{420}$ include the oxidized and reduced forms, total coenzyme B includes HS-CoB and CoM-S-S-CoB, and total MFR includes MFR and CHO-MFR. The model is insensitive to initial concentrations of non-coupled metabolites.

**Table S4:** Data for isotopic model calibration. For a complete list of the data used see the SM. The culture conditions are abbreviated for pure cultures (PC), co-cultures (CC) and enrichment cultures (EC).

| **Source** | $^{13}\alpha_{CO_2-CH_4}$ | $^{2}\alpha_{CH_4-H_2O}$ | $\Delta^{13}CH_3D$ | $\Delta^{12}CH_2D_2$ | **Temp. (°C)** | **Culture** | **Ref.** |
|---|---|---|---|---|---|---|---|
| Botz et al., 1996 | + | − | − | − | 35–85 | PC | (14) |
| Valentine et al., 2004 | + | − | − | − | 40–75 | PC | (31) |
| Penning et al., 2005 | + | − | − | − | 30–37 | CC | (2) |
| Yoshioka et al., 2008 | − | + | − | − | 55–65 | CC/PC | (27) |
| Takai et al., 2008 | + | − | − | − | 85–122 | PC | (32) |
| Hattori et al., 2012 | + | + | − | − | 55–65 | EC | (33) |
| Stolper et al., 2014 | − | + | + | − | 30–70 | PC | (72) |
| Wang et al., 2015 | + | + | + | − | 25–80 | PC | (24) |
| Okumura et al., 2016 | + | + | − | − | 25–60 | CC/PC | (3) |
| Young et al., 2017 | + | + | + | + | 55–65 | PC | (73) |
| Gruen et al., 2018 | + | + | + | − | 21–80 | PC | (74) |
| Topçuoğlu et al., 2019 | + | − | − | − | 83 | CC/PC | (34) |

**Table S5:** Ranges of $H_2$ concentrations in natural environments.

| **Environment** | **$H_2$ Range (nM)** | **Source** | **Ref.** |
|---|---|---|---|
| Freshwater sediments | 20 | Conrad et al., 1985<br>Lovley et al., 1988 | (4, 75) |
| Marine sediments | | | |
| *shallow* | 0.5–5 | Novelli et al., 1988<br>Hoehler et al., 1998<br>Zhuang et al., 2018 | (76–78) |
| *deep* | 100 | Ijiri et al., 2018 | (79) |
| Estuaries | 10–100 | Michener et al., 1988 | (80) |
| Rice paddies | 15–60 | Yao et al., 1999 | (81) |

**Table S6:** Estimated environmental csMR based on the relations between measured cell densities and bulk methanogenesis rates in shallow and deep marine sediments.

| Source | Max. sediment depth (m) | Temp. (°C) | Cells | bMR | csMR | Ref. |
|---|---|---|---|---|---|---|
| Hoehler et al., 1994 (W) | 0.4 | 6 | $4.3\times10^{7}$ | $1.1^{\dagger\,\S}$ | $2.6\times10^{-2}$ | (82) |
| Hoehler et al., 1994 (S) | 0.4 | 28 | $4.3\times10^{7}$ | $32^{\dagger\,\S}$ | $7.5\times10^{-1}$ | (82) |
| Claypool et al., 2006 | 120 | 10 | $2.9\times10^{5}$ | $7.2\times10^{-4}$ | $2.4\times10^{-3}$ | (83) |
| Sivan et al., 2007 | 200 | 10 | $1.3\times10^{5}$ | $2.3\times10^{-4}$ | $1.8\times10^{-3}$ | (84) |
| Parkes et al., 2007 | 4 | 16 | $5.4\times10^{7\P}$ | $4.7\times10^{-1\dagger}$ | $8.7\times10^{-3}$ | (85) |
| Beulig et al., 2018 | 1.4 | 8 | $9.5\times10^{5\P}$ | $4.8\times10^{-1\dagger}$ | $5.0\times10^{-1}$ | (86) |
| Chuang et al., 2018 | 40 | 5 | $6.8\times10^{5}$ | $4.1\times10^{-4}$ | $6.0\times10^{-4}$ | (87) |
| Zhuang et al., 2018‡ | 6 | 13 | $7.9\times10^{6}$ | $7.6\times10^{-1}$ | $9.5\times10^{-2}$ | (78) |
| Zhuang et al., 2018$ | 5 | 13 | $7.9\times10^{6}$ | $1.0\times10^{-3}$ | $1.3\times10^{-4}$ | (78) |

Where not measured directly, cell densities were estimated based on their dependence on depth (28, 29) and assuming that of these cells 12% are Archaea in open-ocean sites and 40% in ocean margin sites (30), and that 50% of Archaea are methanogens (29).

† $^{14}C$ tracer measurements.

§ Tracer measurements were done for both gross methane production and consumption, and the results are shown as a net rate, which is calculated as the difference between the measured forward and backward rates.

¶ Direct cell counts.

‡ Rhone River pro-delta.

$ Gulf of Lion shelf.

### References


[1] A. Missner, P. Kügler, S. M. Saparov, K. Sommer, J. C. Mathai, M. L. Zeidel, P. Pohl, Carbon dioxide transport through membranes. *J. Biol. Chem.* **283**, 25340–7 (2008).

[2] H. Penning, C. M. Plugge, P. E. Galand, R. Conrad, Variation of carbon isotope fractionation in hydrogenotrophic methanogenic microbial cultures and environmental samples at different energy status. *Glob. Change Biol.* **11**, 2103–2113 (2005).

[3] T. Okumura, S. Kawagucci, Y. Saito, Y. Matsui, K. Takai, H. Imachi, Hydrogen and carbon isotope systematics in hydrogenotrophic methanogenesis under H2-limited and H2-enriched conditions: Implications for the origin of methane and its isotopic diagnosis. *Prog. Earth Planet. Sci.* **3**, 14 (2016).

[4] R. Conrad, T. J. Phelps, J. G. Zeikus, Gas metabolism evidence in support of the juxtaposition of hydrogen-producing and methanogenic bacteria in sewage sludge and lake sediments. *Appl. Environ. Microbiol.* **50**, 595–601 (1985).

[5] E. Giraldo-Gomez, S. Goodwin, M. S. Switzenbaum, Influence of mass transfer limitations on determination of the half saturation constant for hydrogen uptake in a mixed-culture CH4-producing enrichment. *Biotechnol. Bioeng.* **40**, 768–76 (1992).

[6] T. Lienard, B. Becher, M. Marschall, S. Bowien, G. Gottschalk, Sodium Ion Translocation by N5-Methyltetrahydromethanopterin: Coenzyme M Methyltransferase from Methanosarcina mazei Go1 Reconstituted in Ether Lipid Liposomes. *Eur. J. Biochem.* **239**, 857–864 (1996).

[7] U. Deppenmeier, V. Müller, Life close to the thermodynamic limit: How methanogenic archaea conserve energy. *Results Probl. Cell Differ.* **45**, 123–52 (2008).

[8] R. K. Thauer, S. Shima, *Annals of the New York Academy of Sciences* (2008), vol. 1125, pp. 158–170.

[9] A. Flamholz, E. Noor, A. Bar-Even, R. Milo, eQuilibrator–the biochemical thermodynamics calculator. *Nucleic Acids Res.* **40**, D770-5 (2012).

[10] A. Bar-Even, E. Noor, Y. Savir, W. Liebermeister, D. Davidi, D. S. Tawfik, R. Milo, The Moderately Efficient Enzyme: Evolutionary and Physicochemical Trends Shaping Enzyme Parameters. *Biochemistry* **50**, 4402–4410 (2011).

[11] J. L. Pennings, P. Vermeij, L. M. de Poorter, J. T. Keltjens, G. D. Vogels, Adaptation of methane formation and enzyme contents during growth of Methanobacterium thermoautotrophicum (strain ΔH) in a fed-batch fermentor. *Antonie Van Leeuwenhoek* **77**, 281–291 (2000).

[12] M. Lupascu, J. L. Wadham, E. R. C. Hornibrook, R. D. Pancost, Temperature Sensitivity of Methane Production in the Permafrost Active Layer at Stordalen, Sweden: A Comparison with Non-permafrost Northern Wetlands. *Arct. Antarct. Alp. Res.* **44**, 469–482 (2012).

[13] Q. Wu, R. Ye, S. D. Bridgham, Q. Jin, Limitations of the Q10 Coefficient for Quantifying Temperature Sensitivity of Anaerobic Organic Matter Decomposition: A Modeling Based Assessment. *J. Geophys. Res. Biogeosciences* **126**, e2021JG006264 (2021).

[14] R. Botz, H. D. Pokojski, M. Schmitt, M. Thomm, Carbon isotope fractionation during bacterial methanogenesis by CO2 reduction. *Org. Geochem.* **25**, 255–262 (1996).

[15] P. Vermeij, J. L. Pennings, S. M. Maassen, J. T. Keltjens, G. D. Vogels, Cellular levels of factor 390 and methanogenic enzymes during growth of Methanobacterium thermoautotrophicum deltaH. *J. Bacteriol.* **179**, 6640–8 (1997).

[16] S. Kato, T. Kosaka, K. Watanabe, Comparative transcriptome analysis of responses of Methanothermobacter thermautotrophicus to different environmental stimuli. *Environ. Microbiol.* **10**, 893–905 (2008).

[17] Q. Xia, T. Wang, E. L. Hendrickson, T. J. Lie, M. Hackett, J. A. Leigh, Quantitative proteomics of nutrient limitation in the hydrogenotrophic methanogen Methanococcus maripaludis. *BMC Microbiol.* **9**, 149 (2009).

[18] J. Gropp, M. A. Iron, I. Halevy, Theoretical estimates of equilibrium carbon and hydrogen isotope effects in microbial methane production and anaerobic oxidation of methane. *Geochimica et Cosmochimica Acta* **295**, 237–264 (2021).

[19] J. Gropp, M. A. Iron, I. Halevy, Corrigendum to "Theoretical estimates of equilibrium carbon and hydrogen isotope effects in microbial methane production and anaerobic oxidation of methane" [Geochim. Cosmochim. Acta 295 (2021) 237–264]. *Geochimica et Cosmochimica Acta* **306**, 386–389 (2021).

[20] S. Scheller, M. Goenrich, R. K. Thauer, B. Jaun, Methyl-coenzyme M reductase from methanogenic archaea: Isotope effects on the formation and anaerobic oxidation of methane. *J. Am. Chem. Soc.* **135**, 14975–84 (2013).

[21] B. Fry, H. Gest, J. Hayes, Isotope effects associated with the anaerobic oxidation of sulfite and thiosulfate by the photosynthetic bacterium, Chromatium vinosum. *FEMS Microbiology Letters* **27**, 227–232 (1985).

[22] A. Sattler, Hydrogen/Deuterium (H/D) Exchange Catalysis in Alkanes. *ACS Catal.* **8**, 2296–2312 (2018).

[23] M. Gómez-Gallego, M. A. Sierra, Kinetic isotope effects in the study of organometallic reaction mechanisms. *Chem. Rev.* **111**, 4857–963 (2011).

[24] D. T. Wang, D. S. Gruen, B. S. Lollar, K.-U. Hinrichs, L. C. Stewart, J. F. Holden, A. N. Hristov, J. W. Pohlman, P. L. Morrill, M. Könneke, K. B. Delwiche, E. P. Reeves, C. N. Sutcliffe, D. J. Ritter, J. S. Seewald, J. C. McIntosh, H. F. Hemond, M. D. Kubo, D. Cardace, T. M. Hoehler, S. Ono, Nonequilibrium clumped isotope signals in microbial methane. *Science* **348**, 428–431 (2015).

[25] D. T. Wang, P. V. Welander, S. Ono, Fractionation of the methane isotopologues 13CH4, 12CH3D, and 13CH3D during aerobic oxidation of methane by Methylococcus capsulatus (Bath). *Geochim. Cosmochim. Acta* **192**, 186–202 (2016).

[26] X. Cao, H. Bao, Y. Peng, A kinetic model for isotopologue signatures of methane generated by biotic and abiotic CO2 methanation. *Geochim. Cosmochim. Acta* **249**, 59–75 (2019).

[27] H. Yoshioka, S. Sakata, Y. Kamagata, Hydrogen isotope fractionation by Methanothermobacter thermoautotrophicus in coculture and pure culture conditions. *Geochim. Cosmochim. Acta* **72**, 2687–2694 (2008).

[28] J. Kallmeyer, R. Pockalny, R. R. Adhikari, D. C. Smith, S. D'Hondt, Global distribution of microbial abundance and biomass in subseafloor sediment. *Proc. Natl. Acad. Sci.* **109**, 16213–16216 (2012).

[29] C. Petro, P. Starnawski, A. Schramm, K. U. Kjeldsen, Microbial community assembly in marine sediments. *Aquat. Microb. Ecol.* **79**, 177–195 (2017).

[30] T. Hoshino, F. Inagaki, Abundance and distribution of Archaea in the subseafloor sedimentary biosphere. *ISME J.* **13**, 227–231 (2019).

[31] D. L. Valentine, A. Chidthaisong, A. Rice, W. S. Reeburgh, S. C. Tyler, Carbon and hydrogen isotope fractionation by moderately thermophilic methanogens. *Geochim. Cosmochim. Acta* **68**, 1571–1590 (2004).

[32] K. Takai, K. Nakamura, T. Toki, U. Tsunogai, M. Miyazaki, J. Miyazaki, H. Hirayama, S. Nakagawa, T. Nunoura, K. Horikoshi, Cell proliferation at 122 degrees C and isotopically heavy CH4 production by a hyperthermophilic methanogen under high-pressure cultivation. *Proc. Natl. Acad. Sci. U. S. A.* **105**, 10949–54 (2008).

[33] S. Hattori, H. Nashimoto, H. Kimura, K. Koba, K. Yamada, M. Shimizu, H. Watanabe, M. Yoh, N. Yoshida, Hydrogen and carbon isotope fractionation by thermophilic hydrogenotrophic methanogens from a deep aquifer under coculture with fermenters. *Geochem. J.* **46**, 193–200 (2012).

[34] B. D. Topçuoğlu, C. Meydan, T. B. Nguyen, S. Q. Lang, J. F. Holden, Growth Kinetics, Carbon Isotope Fractionation, and Gene Expression in the Hyperthermophile Methanocaldococcus jannaschii during Hydrogen-Limited Growth and Interspecies Hydrogen Transfer. *Appl. Environ. Microbiol.* **85**, 1–14 (2019).

[35] R. A. Alberty, *Thermodynamics of Biochemical Reactions* (John Wiley & Sons, Inc., Hoboken, NJ, USA, 2003).

[36] P. A. Bertram, R. K. Thauer, Thermodynamics of the Formylmethanofuran Dehydrogenase Reaction in Methanobacterium Thermoautotrophicum. *Eur. J. Biochem.* **226**, 811–818 (1994).

[37] W. Buckel, R. K. Thauer, Energy conservation via electron bifurcating ferredoxin reduction and proton/Na(+) translocating ferredoxin oxidation. *Biochim. Biophys. Acta* **1827**, 94–113 (2013).

[38] B. E. Maden, Tetrahydrofolate and tetrahydromethanopterin compared: Functionally distinct carriers in C1 metabolism. *Biochem. J.* **350 Pt 3**, 609–29 (2000).

[39] F. Jacobson, C. Walsh, Properties of 7,8-Didemethyl-8-hydroxy-5-deazaflavins Relevant to Redox Coenzyme Function in Methanogen Metabolism. *Biochemistry* **23**, 979–988 (1984).

[40] M. Tietze, A. Beuchle, I. Lamla, N. Orth, M. Dehler, G. Greiner, U. Beifuss, Redox Potentials of Methanophenazine and CoB-S-S-CoM, Factors Involved in Electron Transport in Methanogenic Archaea. *ChemBioChem* **4**, 333–335 (2003).

[41] G. Börner, M. Karrasch, R. K. Thauer, Molybdopterin adenine dinucleotide and molybdopterin hypoxanthine dinucleotide in formylmethanofuran dehydrogenase from Methanobacterium thermoautotrophicum (Marburg). *FEBS Lett.* **290**, 31–4 (1991).

[42] P. A. Bertram, M. Karrasch, R. A. Schmitz, R. Bocher, S. P. J. Albracht, R. K. Thauer, Formylmethanofuran dehydrogenases from methanogenic Archaea Substrate specificity, EPR properties and reversible inactivation by cyanide of the molybdenum or tungsten iron-sulfur proteins. *Eur. J. Biochem.* **220**, 477–484 (1994).

[43] M. I. Donnelly, R. S. Wolfe, The role of formylmethanofuran: Tetrahydromethanopterin formyltransferase in methanogenesis from carbon dioxide. *J. Biol. Chem.* **261**, 16653–9 (1986).

[44] J. Breitung, R. K. Thauer, Formylmethanofuran: Tetrahydromethanopterin formyltransferase from Methanosarcina barkeri. Identification of N5-formyltetrahydromethanopterin as the product. *FEBS Lett.* **275**, 226–30 (1990).

[45] A. A. DiMarco, M. I. Donnelly, R. S. Wolfe, Purification and properties of the 5,10-methenyltetrahydromethanopterin cyclohydrolase from Methanobacterium thermoautotrophicum. *J. Bacteriol.* **168**, 1372–7 (1986).

[46] J. Breitung, R. A. Schmitz, K. O. Stetter, R. K. Thauer, N5,N10-Methenyltetrahydromethanopterin cyclohydrolase from the extreme thermophile Methanopyrus kandleri: Increase of catalytic efficiency (kcat/KM) and thermostability in the presence of salts. *Arch. Microbiol.* **156**, 517–524 (1991).

[47] B. W. te Brömmelstroet, C. M. Hensgens, W. J. Geerts, J. T. Keltjens, C. van der Drift, G. D. Vogels, Purification and properties of 5,10-methenyltetrahydromethanopterin cyclohydrolase from Methanosarcina barkeri. *J. Bacteriol.* **172**, 564–71 (1990).

[48] B. Mukhopadhyay, L. Daniels, Aerobic purification of N5,N10-methylenetetrahydromethanopterin dehydrogenase, separated from N5,N10-methenyltetrahydromethanopterin cyclohydrolase, from Methanobacterium thermoautotrophicum strain Marburg. *Can. J. Microbiol.* **35**, 499–507 (1989).

[49] B. te Brömmelstroet, C. M. Hensgens, J. T. Keltjens, C. van der Drift, G. D. Vogels, Purification and characterization of coenzyme F420-dependent 5,10-methylenetetrahydromethanopterin dehydrogenase from Methanobacterium thermoautotrophicum strain ΔH. *Biochim. Biophys. Acta BBA - Gen. Subj.* **1073**, 77–84 (1991).

[50] C. Zirngibl, W. Dongen, B. Schworer, R. Bunau, M. Richter, A. Klein, R. K. Thauer, H2-forming methylenetetrahydromethanopterin dehydrogenase, a novel type of hydrogenase without iron-sulfur clusters in methanogenic archaea. *Eur. J. Biochem.* **208**, 511–520 (1992).

[51] E. J. Lyon, S. Shima, G. Buurman, S. Chowdhuri, A. Batschauer, K. Steinbach, R. K. Thauer, UV-A/blue-light inactivation of the 'metal-free' hydrogenase (Hmd) from methanogenic archaea. *Eur. J. Biochem.* **271**, 195–204 (2004).

[52] G. C. Hartmann, A. R. Klein, M. Linder, R. K. Thauer, Purification, properties and primary structure of H2-formingN5,N10-methylenetetrahydromethanopterin dehydrogenase from Methanococcus thermolithotrophicus. *Arch. Microbiol.* **165**, 187–193 (1996).

[53] K. Ma, R. K. Thauer, Purification and properties of N5, N10-methylenetetrahydromethanopterin reductase from Methanobacterium thermoautotrophicum (strain Marburg). *Eur. J. Biochem.* **191**, 187–193 (1990).

[54] B. W. te Brömmelstroet, W. J. Geerts, J. T. Keltjens, C. van der Drift, G. D. Vogels, C. van der Drift, G. D. Vogels, Purification and properties of 5,10-methylenetetrahydromethanopterin dehydrogenase and 5,10-methylenetetrahydromethanopterin reductase, two coenzyme F420-dependent enzymes, from Methanosarcina barkeri. *Biochim. Biophys. Acta BBA - Protein Struct. Mol. Enzymol.* **1079**, 293–302 (1991).

[55] V. R. Vepachedu, J. G. Ferry, Role of the Fused Corrinoid/Methyl Transfer Protein CmtA during CO-Dependent Growth of Methanosarcina acetivorans. *J. Bacteriol.* **194**, 4161–4168 (2012).

[56] S. Rospert, R. Böcher, S. Albracht, R. Thauer, Methyl-coenzyme M reductase preparations with high specific activity from H2-preincubated cells of Methanobacterium thermoautotrophicum. *FEBS Lett.* **291**, 371–375 (1991).

[57] L. G. Bonacker, S. Baudner, E. Mörschel, R. Böcher, R. K. Thauer, Properties of the two isoenzymes of methyl-coenzyme M reductase in Methanobacterium thermoautotrophicum. *Eur. J. Biochem.* **217**, 587–95 (1993).

[58] M. Dey, X. Li, R. C. Kunz, S. W. Ragsdale, Detection of Organometallic and Radical Intermediates in the Catalytic Mechanism of Methyl-Coenzyme M Reductase Using the Natural Substrate Methyl-Coenzyme M and a Coenzyme B Substrate Analogue. *Biochemistry* **49**, 10902–10911 (2010).

[59] P. E. Rouvière, T. A. Bobik, R. S. Wolfe, Reductive activation of the methyl coenzyme M methylreductase system of Methanobacterium thermoautotrophicum delta H. *J. Bacteriol.* **170**, 3946–52 (1988).

[60] J. Ellermann, S. Rospert, R. K. Thauer, M. Bokranz, A. Klein, M. Voges, A. Berkessel, Methylcoenzyme-M reductase from Methanobacterium thermoautotrophicum (strain Marburg). Purity, activity and novel inhibitors. *Eur. J. Biochem.* **184**, 63–68 (1989).

[61] J. A. Fox, D. J. Livingston, W. H. Orme-Johnson, C. T. Walsh, 8-Hydroxy-5-deazaflavin-reducing hydrogenase from Methanobacterium thermoautotrophicum: 1. Purification and characterization. *Biochemistry* **26**, 4219–4227 (1987).

[62] A.-K. Kaster, J. Moll, K. Parey, R. K. Thauer, Coupling of ferredoxin and heterodisulfide reduction via electron bifurcation in hydrogenotrophic methanogenic archaea. *PNAS* **108**, 2981–6 (2011).

[63] E. Setzke, R. Hedderich, S. Heiden, R. K. Thauer, H2: Heterodisulfide oxidoreductase complex from Methanobacterium thermoautotrophicum. Composition and properties. *Eur. J. Biochem. FEBS* **220**, 139–48 (1994).

[64] J. Meuer, S. Bartoschek, J. Koch, A. Kunkel, R. Hedderich, Purification and catalytic properties of Ech hydrogenase from Methanosarcina barkeri. *Eur. J. Biochem.* **265**, 325–335 (1999).

[65] R. Hedderich, A. Berkessel, R. K. Thauer, Purification and properties of heterodisulfide reductase from Methanobacterium thermoautotrophicum (strain Marburg). *Eur. J. Biochem.* **193**, 255–261 (1990).

[66] P. Schönheit, H. J. Perski, ATP synthesis driven by a potassium diffusion potential in Methanobacterium thermoautotrophicum is stimulated by sodium. *FEMS Microbiol. Lett.* **20**, 263–267 (1983).

[67] M. A. Lever, K. L. Rogers, K. G. Lloyd, J. Overmann, B. Schink, R. K. Thauer, T. M. Hoehler, B. B. Jørgensen, Life under extreme energy limitation: A synthesis of laboratory- and field-based investigations. *FEMS Microbiol Rev* **39**, 688–728 (2015).

[68] E. Heine-Dobbernack, S. M. Schoberth, H. Sahm, Relationship of Intracellular Coenzyme F(420) Content to Growth and Metabolic Activity of Methanobacterium bryantii and Methanosarcina barkeri. *Appl. Environ. Microbiol.* **54**, 454–9 (1988).

[69] R. Sander, Compilation of Henry's law constants (version 4.0) for water as solvent. *Atmospheric Chem. Phys.* **15**, 4399–4981 (2015).

[70] A. M. Feist, J. C. M. Scholten, B. Ø. Palsson, F. J. Brockman, T. Ideker, Modeling methanogenesis with a genome-scale metabolic reconstruction of Methanosarcina barkeri. *Mol. Syst. Biol.* **2**, 2006.0004 (2006).

[71] L. M. I. de Poorter, W. G. Geerts, A. P. R. Theuvenet, J. T. Keltjens, Bioenergetics of the formyl-methanofuran dehydrogenase and heterodisulfide reductase reactions in Methanothermobacter thermautotrophicus. *Eur. J. Biochem.* **270**, 66–75 (2002).

[72] D. A. Stolper, M. Lawson, C. L. Davis, A. A. Ferreira, E. V. Santos Neto, G. S. Ellis, M. D. Lewan, A. M. Martini, Y. Tang, M. Schoell, A. L. Sessions, J. M. Eiler, Formation temperatures of thermogenic and biogenic methane. *Science* **344**, 1500–1503 (2014).

[73] E. D. Young, I. Kohl, B. S. Lollar, G. Etiope, D. Rumble, S. Li, M. Haghnegahdar, E. Schauble, K. McCain, D. Foustoukos, C. Sutclife, O. Warr, C. Ballentine, T. Onstott, H. Hosgormez, A. Neubeck, J. Marques, I. Pérez-Rodríguez, A. Rowe, D. LaRowe, C. Magnabosco, L. Yeung, J. Ash, L. Bryndzia, The relative abundances of resolved 12CH2D2 and 13CH3D and mechanisms controlling isotopic bond ordering in abiotic and biotic methane gases. *Geochim. Cosmochim. Acta* **203**, 235–264 (2017).

[74] D. S. Gruen, D. T. Wang, M. Könneke, B. D. Topçuoğlu, L. C. Stewart, T. Goldhammer, J. F. Holden, K.-U. Hinrichs, S. Ono, Experimental investigation on the controls of clumped isotopologue and hydrogen isotope ratios in microbial methane. *Geochim. Cosmochim. Acta* **237**, 339–356 (2018).

[75] D. R. Lovley, S. Goodwin, Hydrogen concentrations as an indicator of the predominant terminal electron-accepting reactions in aquatic sediments. *Geochim. Cosmochim. Acta* **52**, 2993–3003 (1988).

[76] P. C. Novelli, A. R. Michelson, M. I. Scranton, G. T. Banta, J. E. Hobbie, R. W. Howarth, Hydrogen and acetate cycling in two sulfate-reducing sediments: Buzzards Bay and Town Cove, Mass. *Geochimica et Cosmochimica Acta* **52**, 2477–2486 (1988).

[77] T. M. Hoehler, M. J. Alperin, D. B. Albert, C. S. Martens, Thermodynamic control on hydrogen concentrations in anoxic sediments. *Geochim. Cosmochim. Acta* **62**, 1745–1756 (1998).

[78] G.-C. Zhuang, V. B. Heuer, C. S. Lazar, T. Goldhammer, J. Wendt, V. A. Samarkin, M. Elvert, A. P. Teske, S. B. Joye, K.-U. Hinrichs, Relative importance of methylotrophic methanogenesis in sediments of the Western Mediterranean Sea. *Geochim. Cosmochim. Acta* **224**, 171–186 (2018).

[79] A. Ijiri, F. Inagaki, Y. Kubo, R. R. Adhikari, S. Hattori, T. Hoshino, H. Imachi, S. Kawagucci, Y. Morono, Y. Ohtomo, S. Ono, S. Sakai, K. Takai, T. Toki, D. T. Wang, M. Y. Yoshinaga, G. L. Arnold, J. Ashi, D. H. Case, T. Feseker, K.-U. Hinrichs, Y. Ikegawa, M. Ikehara, J. Kallmeyer, H. Kumagai, M. A. Lever, S. Morita, K.-i. Nakamura, Y. Nakamura, M. Nishizawa, V. J. Orphan, H. Røy, F. Schmidt, A. Tani, W. Tanikawa, T. Terada, H. Tomaru, T. Tsuji, U. Tsunogai, Y. T. Yamaguchi, N. Yoshida, Deep-biosphere methane production stimulated by geofluids in the Nankai accretionary complex. *Sci. Adv.* **4**, eaao4631 (2018).

[80] R. H. Michener, M. I. Scranton, P. Novelli, Hydrogen (H2) distributions in the carmans river estuary. *Estuarine, Coastal and Shelf Science* **27**, 223–235 (1988).

[81] H. Yao, R. Conrad, Thermodynamics of methane production in different rice paddy soils from China, the Philippines and Italy. *Soil Biology and Biochemistry* **31**, 463–473 (1999).

[82] T. M. Hoehler, M. J. Alperin, D. B. Albert, C. S. Martens, Field and laboratory studies of methane oxidation in an anoxic marine sediment: Evidence for a methanogen-sulfate reducer consortium. *Glob. Biogeochem. Cycles* **8**, 451–463 (1994).

[83] G. E. Claypool, A. V. Milkov, Y. J. Lee, M. E. Torres, W. S. Borowski, H. Tomaru, Microbial methane generation and gas transport in shallow sediments of an accretionary complex, southern hydrate ridge (ODP Leg 204), offshore Oregon, USA. *Proc. Ocean Drill. Program Sci. Results* **204** (2006).

[84] O. Sivan, D. P. Schrag, R. W. Murray, Rates of methanogenesis and methanotrophy in deep-sea sediments. *Geobiology* **5**, 141–151 (2007).

[85] R. J. Parkes, B. A. Cragg, N. Banning, F. Brock, G. Webster, J. C. Fry, E. Hornibrook, R. D. Pancost, S. Kelly, N. Knab, B. B. Jørgensen, J. Rinna, A. J. Weightman, Biogeochemistry and biodiversity of methane cycling in subsurface marine sediments (Skagerrak, Denmark). *Environ. Microbiol.* **9**, 1146–1161 (2007).

[86] F. Beulig, H. Røy, S. E. McGlynn, B. B. Jørgensen, Cryptic CH4 cycling in the sulfate–methane transition of marine sediments apparently mediated by ANME-1 archaea. *ISME J.* **13**, 250–262 (2018).

[87] P.-C. Chuang, T. Frank Yang, K. Wallmann, R. Matsumoto, C.-Y. Hu, H.-W. Chen, S. Lin, C.-H. Sun, H.-C. Li, Y. Wang, A. W. Dale, Carbon isotope exchange during anaerobic oxidation of methane (AOM) in sediments of the northeastern South China Sea. *Geochim. Cosmochim. Acta* **246**, 138–155 (2018).
